# Metabolism-dependent secondary effect of anti-MAPK cancer therapy on DNA repair

**DOI:** 10.1101/2023.06.19.544800

**Authors:** Fabien Aubé, Nicolas Fontrodona, Laura Guiguettaz, Elodie Vallin, Audrey Lapendry, Emiliano P. Ricci, Didier Auboeuf

**Affiliations:** Laboratoire de Biologie et Modélisation de la Cellule, Ecole Normale Supérieure de Lyon, CNRS, UMR 5239, Inserm, U1293, Université Claude Bernard Lyon 1, 46 allée d’Italie F-69364 Lyon, France; Equipe Labellisée Ligue Nationale Contre le Cancer, LBMC, ENS Lyon

## Abstract

Amino acid bioavailability impacts mRNA translation in a codon depending manner. Here, we report that the anti-cancer MAPK inhibitors (MAPKi) decrease the intracellular concentration of aspartate and glutamate in melanoma cells. This results in the accumulation of ribosomes on codons corresponding to these amino acids and triggers the translation-dependent degradation of mRNAs encoding aspartate- and glutamate-rich proteins mostly involved in DNA metabolism. Consequently, cells that survive to MAPKi degrade aspartate and glutamate to generate energy, which simultaneously decreases their needs in amino acids owing to the downregulation of aspartate- and glutamate-rich proteins involved in cell proliferation. Concomitantly, the downregulation of aspartate- and glutamate-rich proteins involved in DNA repair increases DNA damage loads. Thus, DNA repair defects, and therefore mutations, are, at least in part, a secondary effect of the metabolic adaptation of cells exposed to MAPKi.

## Introduction

Some amino acids correspond to several codons, bringing a redundancy into the genetic code, which plays a role in gene product expression level. Indeed, the nature of mRNA codons affects the dynamics of ribosomes, through the time required to "decode" each codon, with consequences on translation efficiency, protein folding and mRNA co-translational degradation through the translation-dependent mRNA decay (TDD) pathway^1–7^. For instance, sub-optimal codons or clusters of rare codons modulate gene product expression levels in a gene- and species-selective manner^1, 3, 4, 7–9^. At least in humans, sub-optimal and rare codons are more frequently A/T-ending codons, while optimal codons are more frequently G/C-ending codons likely because of the more unstable interactions between A/T-ending codons with their cognate anti-codons as compared to G/C-ending codons^4, 10–14^.

While codon usage is an intrinsic parameter of gene products with respect to their translatability and stability, there are many extrinsic parameters that modulate codon-depending effects in a cell type- and context-depending manner. Among the extrinsic parameters, enzymatic-dependent biochemical modifications of anti-codons have been reported to modulate the interactions between codons and anti-codons^15, 16^. For example, the cytosolic thiouridylase 2 (CTU2) enzyme that biochemically modifies the tRNA wobble uridine ─ thereby affecting the decoding of some A-ending codons, such as the AAA and GAA codons ─ is required for the efficient translation of a subset of mRNAs that promote survival and resistance to therapy of BRAF-mutated melanoma cells^17, 18^. In addition, variation in the expression levels of different classes of tRNAs can change the codon-dependent effect on gene product expression level in a cell type- and context-dependent manner^19, 20^. For instance, proliferative cells express higher levels of tRNAs corresponding to A/T-ending codons than differentiated cells, leading to the differential expression levels of gene products enriched in either A/T- or G/C-ending codons^21, 22^. Along the same line, the expression levels of aminoacyl transferases, which load amino acids onto tRNAs, can modulate the codon-dependent effects on gene product expression levels^23, 24^. An example is given by the leucyl-tRNA synthetase (LARS), which increases the selective loading of tRNA-Leu^CAG^ isoacceptor and thereby affects the translatability of mRNAs containing the CAG codon^25^.

The bioavailability of amino acids is of particular interest as an extrinsic parameter that modulates codon-dependent effects on gene product expression levels. Indeed, the dynamics of ribosomes depend in part on the intracellular concentration of loaded-tRNAs, which itself depends on the intracellular concentration of amino acids. Thus, the translation of mRNAs requiring an amino acid whose bioavailability decreases can be impacted as shown for numerous amino acids, including non-essential ones such as glutamine^5, 7, 26–29^. As a consequence of the link between amino acid bioavailability and translation, the synthesis of proteins with amino acid composition biases depend on the cell metabolism^23, 25, 28, 30, 31^. For example, a large amount of proline can be produced only under certain metabolic conditions, which therefore determines whether some proline-rich proteins of the extracellular matrix (e.g., collagen) are produced^29, 32, 33^. This illustrates how the cellular metabolism ─ through amino acid bioavailability ─ is coupled to the nature of cell-expressed proteins on which the cell phenotype depends.

The coupling between cell metabolism and the protein-dependent cell phenotype can be illustrated by the competition between energy production and gene product biogenesis on which depends cell proliferation because some amino acids like glutamine (Gln), glutamate (Glu) and aspartate (Asp) are at the crossroads between several metabolic pathways and the gene expression process. Indeed, the carbon skeleton of these amino acids can either be degraded and end up in the production of energy, or "recycled" to synthesize other amino acids and nucleotides (and therefore gene products)^34–39^. As a consequence, some amino acids may either be used by the cells to produce energy through their complete degradation or be used for the synthesis of large amount of gene products as during cell proliferation. This may explain why such amino acids play a particularly important role in cancer cells, which have a high proliferation rate that requires gene product synthesis, but are in a resource-impoverished micro-environment as a consequence of cell proliferation^35, 37, 40–45^. Accordingly, cancers cells are often addicted to certain amino acids, such as Gln in melanoma cells^37, 41, 44, 45^. The link between cell metabolism and gene expression–dependent cell phenotypes could have consequences in cancer cells exposed to anti-cancer agents, such as MAPK-inhibitors (MAPKi) used to treat melanoma, since these molecules modify the cancer cell metabolism^46–51^. In other words, anticancer therapies may impact cellular phenotypes because of metabolic-dependent effects on gene product expression levels.

Here, we report in a melanoma cell line that MAPKi-downregulated mRNAs encode proteins enriched for certain amino acids, including Glu and Asp whose intracellular concentration decreased in MAPKi-treated cells. Interestingly, MAPKi-downregulated mRNAs encoded proteins involved in cell proliferation and DNA repair, two classes of proteins that are globally enriched in Glu and Asp residues. In line with this observation, MAPKi-treated cells show DNA repair defects. Our results therefore support a model in which the metabolic-dependent effects of MAPKi therapy could result in secondary defects of DNA repair, which could increase the probability of genetically adapted cancer cells to emerge after MAPKi therapy.

## Results

### Compositional biases of MAPKi-regulated gene products

RNA-sequencing was performed after culturing the A375 melanoma cell line for 24h in the absence or presence of a combination of BRAF- and MEK-inhibitors (hereinafter termed MAPK inhibitors [MAPKi]). The expression levels of 2010 or 1719 mRNAs were significantly decreased (downregulated mRNAs) or increased (upregulated mRNAs), respectively, in MAPKi-treated cells as compared to control cells (Fig. 1a and Supplementary Table 1). Interestingly, 813 and 753 MAPKi-downregulated mRNAs encoded proteins associated with the GO terms "nucleoplasm" and "cytoplasm", respectively, and 626 MAPKi-upregulated mRNAs encoded proteins associated with the GO term "Integral component of membrane" (Fig. 1a). In agreement with the fact that nucleoplasmic and cytoplasmic proteins are typically hydrophilic soluble proteins, while membrane proteins tend to be hydrophobic proteins, we noticed that the hydrophobicity index of proteins encoded by upregulated-mRNAs was higher than the hydrophobicity index of proteins encoded by downregulated-mRNAs (Supplementary Fig. 1a). This observation raised the possibility that proteins encoded by down- or up-regulated mRNAs had different amino acid composition biases.

**Figure 1.**
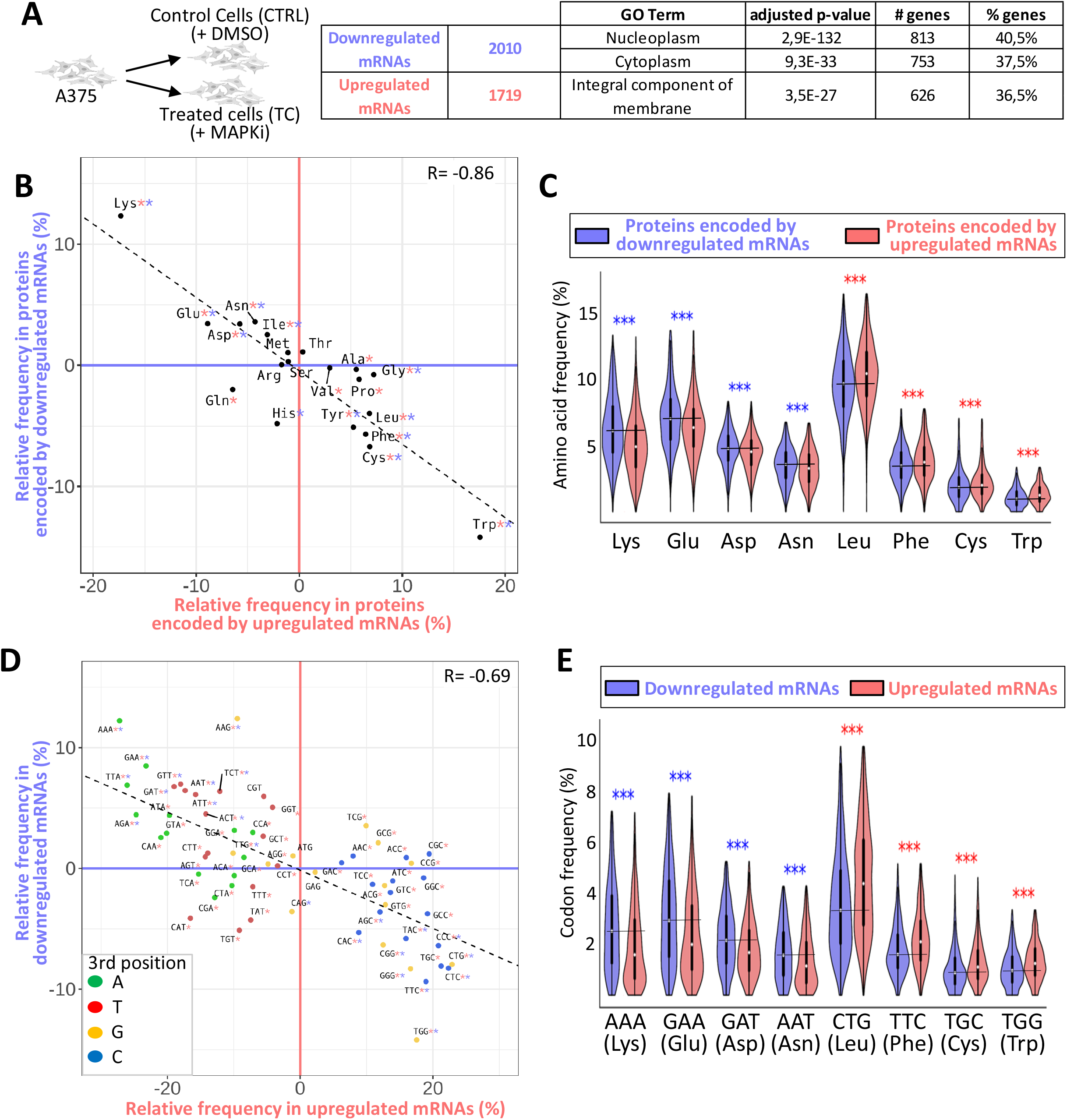
**a** A375 cells were cultured for 24h in the absence (CTRL) or in the presence of MAPKi (TC) before RNA-sequencing. Number and GO term analysis of MAPKi-regulated genes. **b** Amino acid relative frequency (%) in proteins encoded by MAPKi-regulated mRNAs. The x-axis and y-axis correspond to the relative frequency (%) of each amino acid computed from proteins encoded by MAPKi-upregulated and –downregulated mRNAs, respectively in comparison to the amino acid frequency in control proteins, i.e. proteins encoded by mRNAs expressed in the A375 cells. * in red or in blue means that the frequency of an amino acid is statistically different (beta regression analysis followed by a Tukey’s test (pairwise comparison) FDR ≤ 0.05) when comparing control proteins to proteins encoded by MAPKi-upregulated or –downregulated mRNAs, respectively. **c** Amino acid frequency in proteins encoded by MAPKi-upregulated mRNAs (red) or by MAPKi-downregulated mRNAs (blue). ***means that amino acids frequencies are statistically different (beta regression analysis followed by a Tukey’s test (pairwise comparison) FDR <0.001) when comparing proteins encoded by MAPKi-downregulated mRNAs or by MAPKi-upregulated mRNAs. **d** Codon relative frequency (%) in MAPKi-regulated mRNAs. The x-axis and y-axis correspond to the relative frequency (%) of each codon computed in MAPKi-upregulated and -downregulated mRNAs, respectively in comparison to all other mRNAs expressed in A375 cells (control mRNAs). * in red or in blue means that the frequency of a codon is statistically different (beta regression analysis followed by a Tukey’s test (pairwise comparison) FDR ≤ 0.05) when comparing control mRNAs to mRNAs that were upregulated or downregulated, respectively in MAPKi-treated cells. Green, red, orange, and blue dots represent A-, T-, G-, and C-ending codons, respectively. **e** Codon frequency in MAPKi-upregulated mRNAs (red) or in MAPKi-downregulated mRNAs (blue) mRNAs. *** means that codons frequencies are statistically different(beta regression analysis followed by a Tukey’s test (pairwise comparison) FDR <0.001) when comparing MAPKi-downregulated mRNAs and MAPKi-upregulated mRNAs.

Accordingly, hydrophilic residues, such as lysine (Lys), Glu, Asp and asparagine (Asn), were enriched in proteins encoded by MAPKi-downregulated mRNAs, while hydrophobic amino acids, like tryptophan (Trp), cysteine (Cys), phenylalanine (Phe), and leucine (Leu), were enriched in proteins encoded by MAPKi-upregulated mRNA (Fig. 1b, c). Thus, proteins encoded by downregulated mRNAs contained on average 30%, 12%, 8%, and 7% more Lys, Glu, Asp and Asn residues, respectively, than proteins encoded by upregulated mRNAs, while the latter contained on average 24%, 11%, 10%, and 9% more Trp, Cys, Phe and Leu residues, respectively. In addition, a larger part of MAPKi-downregulated mRNAs encoded for proteins with a higher frequency of Lys, Glu, Asp and/or Asn residues, while a larger part of MAPKi-upregulated mRNAs encoded for proteins with a higher frequency of Trp, Cys, Phe and/or Leu residues (Supplementary Fig. 1b).

We next analysed the codon content of MAPKi-regulated mRNAs. Down- and up-regulated mRNAs were enriched for different sets of codons (Fig. 1d). Indeed, A/T-ending codons were enriched in MAPKi-downregulated mRNAs, while G/C-ending codons were enriched in MAPKi-upregulated mRNAs. We also noticed a selective enrichment of a subset of synonymous codons since, for example, only the GAA (but not the GAG), the GAT (but not the GAC), and the AAT (but not the AAC) codons ─ corresponding to Glu, Asp and Asn, respectively ─ were enriched in downregulated mRNAs but reduced in upregulated mRNAs (Fig. 1d, e and Supplementary Fig. 1c).

In summary, mRNAs that were down- or up-regulated by MAPKi treatment contained different codon compositional biases and encoded protein sets with different amino acid compositional biases.

### Compositional biases of TDD-regulated mRNAs in response to MAPKi

We next tested the possibility that MAPKi treatment could affect mRNA stability in a translation-dependent manner. For this, we first measured the TDD index of mRNAs by comparing the expression level of mRNAs in cells treated or not with MAPKi, at initial condition or 3 and 5 hours after inhibition of transcription alone or after inhibition of both transcription and translation (Fig. 2a). MAPKi treatment increased the TDD index of 1390 mRNAs and decreased the TDD index of 183 mRNAs (Fig. 2a and Supplementary Table 1). Although not a formal proof, this observation suggested that MAPKi could affect mRNA expression levels by modulating mRNA stability in a translation-dependent manner.

**Figure 2.**
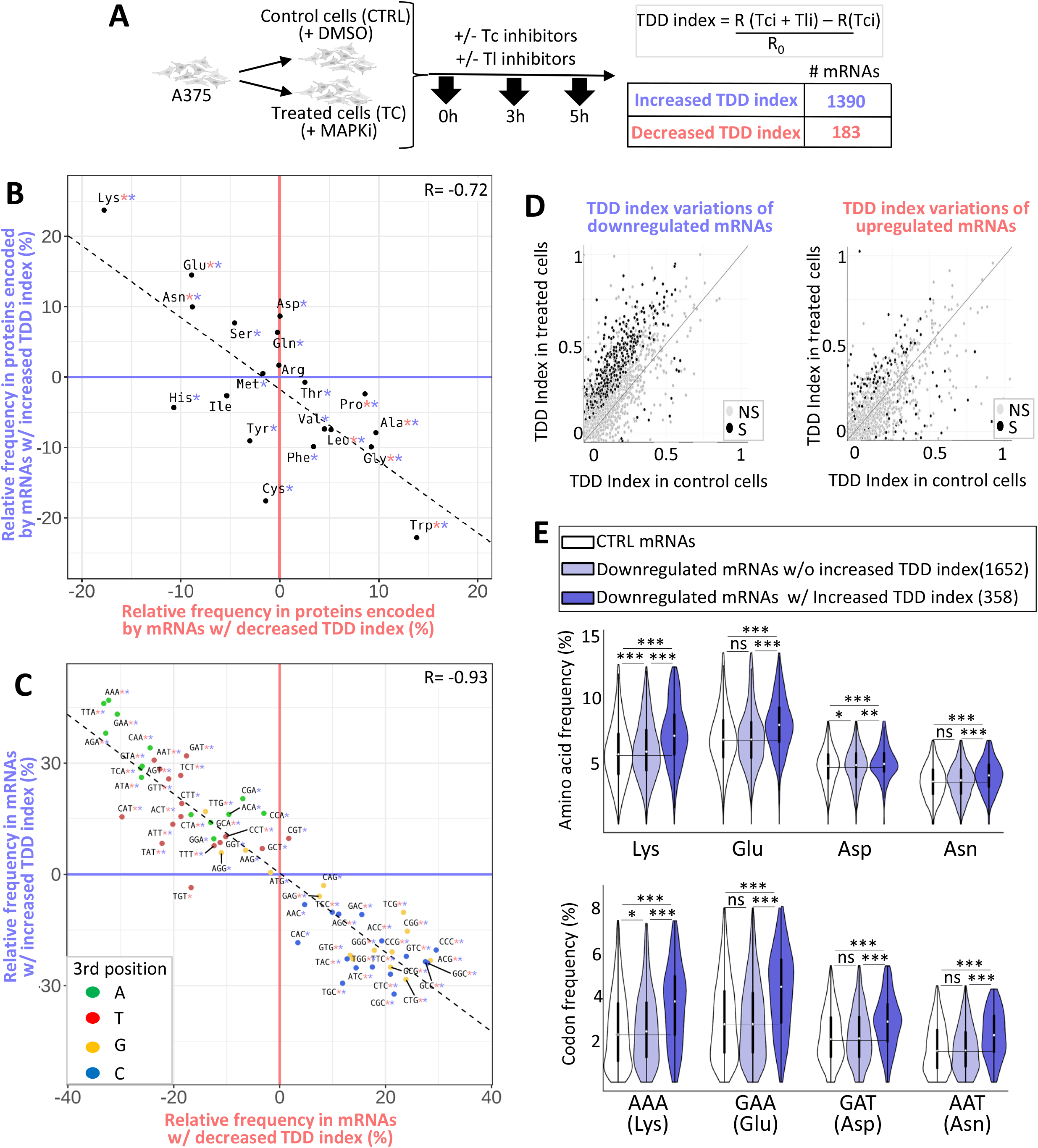
**a** A375 cells were cultured for 18h in the absence or in the presence of MAPKi and were exposed to transcription inhibitors (Tci) and/or to translation inhibitors (Tli) for 0, 3, or 5 h. The TDD index of each A375-expressed mRNA was computed by subtracting the normalized-number of reads (R) obtained in the presence of both Tci and Tli (R(Tci + Tli)) to the normalized-number of reads obtained in the presence of only Tci (R(Tci)). This subtraction was next divided by the initial normalized-number of reads (R0). **b** Amino acid relative frequency (%) in proteins encoded by mRNAs whose TDD was regulated by MAPKi. The x-axis and y-axis correspond to the relative frequency (%) of each amino acid computed from proteins encoded by mRNAs whose TDD index was decreased or increased, respectively in comparison to control proteins, i.e. all other proteins encoded by mRNAs expressed in the A375 cells (control proteins). * in red or in blue means that the frequency of an amino acid is statistically different (beta regression analysis followed by a Tukey’s test (pairwise comparison) FDR ≤ 0.05) when comparing control proteins to proteins encoded by mRNAs whose TDD was decreased or increased, respectively in MAPKi-treated cells. **c** Codon relative frequency (%) in mRNAs whose TDD was regulated by MAPKi. The x-axis and y-axis correspond to the relative frequency (%) of each codon computed from mRNAs whose TDD decreased or increased, respectively in comparison to control mRNAs, i.e., all other mRNAs expressed in A375 cells. * in red or in blue means that the frequency of a codon is statistically different (beta regression analysis followed by a Tukey’s test (pairwise comparison) FDR ≤ 0.05) when comparing control mRNAs to mRNAs whose TDD was decreased or increased, respectively in MAPKi-treated cells. Green, red, orange, and blue dots represent A-, T-, G-, and C-ending codons, respectively. **d** Comparison of the TDD index of mRNAs calculated in control cells or in cells treated for 24h by MAPKi. On the left, the TDD index measured in control cells of each MAPKi-downregulated mRNA (x-axis) was plotted against their TDD index measured in MAPKi-treated cells (y-axis). On the right, the TDD index measured in control cells of each MAPKi-upregulated mRNAs (x-axis) was plotted against their TDD index measured in MAPKi-treated cells (y-axis). Grey dots represent mRNAs whose TDD index was not statistically different (NS) when comparing treated cells to control cells. Black dots represent mRNAs whose TDD index was statistically different (S, linear regression analysis two-tailed t-test p-value ≤ 0.05) when comparing treated cells to control cells. The gray line indicates when the TDD values are identical under the compared conditions. **e** Frequency (%) of codons (on the bottom panel) and amino acids (on the top panel) in three different mRNA populations and the three different protein sets that they produce. On the top, amino acid frequency in proteins encoded by control (CTRL) mRNAs (i.e. expressed mRNAs without those downregulated by MAPKi), MAPKi-downregulated mRNAs whose TDD was not increased, and MAPKi-downregulated mRNAs whose TDD increased. On the bottom, codon frequency in control (CTRL) mRNAs, MAPKi-downregulated mRNAs whose TDD was not increased, and MAPKi-downregulated mRNAs whose TDD increased. *** FDR< 0.001 and * FDR ≤ 0.05 in beta regression analysis followed by a Tukey’s test (pairwise comparison). NS: Not statically significant.

Remarkably, the Lys, Glu, Asp and Asn residues were enriched in proteins encoded by mRNAs with an increased TDD index in response to MAPKi, while other amino acids, including tryptophan (Trp), glycine (Gly), alanine (Ala), proline (Pro) and leucine (Leu), were enriched in proteins encoded by mRNAs with a decreased TDD index in response to MAPKi treatment (Fig. 2b and Supplementary Fig. 2a). In addition, the majority of codons that were enriched in mRNAs with an increased TDD index in response to MAPKi was reduced in mRNAs whose TDD index decreased (Fig. 2c). Furthermore, the majority of codons that was enriched in mRNAs whose TDD index increased in response to MAPKi corresponded to A/T-ending codons, while most codons enriched in mRNAs whose TDD index decreased in response to MAPKi were G/C-ending codons (Fig. 2c and Supplementary Fig. 2b). This observation agreed with a recent report showing that the MAPK pathway modulates codon optimality of A/T-ending codons^8^.

We noticed that MAPKi treatment increased the TDD index of a large number of mRNAs when compared to the number of mRNAs whose TDD decreased (i.e., 1390 vs. 183, Fig. 2a). In addition, a large number of MAPKi-downregulated mRNAs had a significantly increased TDD index in response to MAPKi, while the TDD index of MAPKi-upregulated mRNAs was either increased or decreased (Fig. 2d). This suggested that the TDD increase could contribute to the downregulation of a large subset of mRNAs in response to MAPKi. On the contrary, TDD does not appear to explain MAPKi-induced mRNA upregulation. Based on these considerations, we decided to focus our analyses on MAPKi-downregulated mRNAs whose TDD index increased in response to MAPKi.

Proteins encoded by MAPKi-downregulated mRNAs whose TDD increased had a higher frequency of Lys, Glu, Asp and Asn as compared to proteins encoded by MAPKi-downregulated mRNAs whose TDD was not affected by MAPKi (Fig. 2e, upper panel). Likewise, MAPKi-downregulated mRNAs whose TDD increased had a higher frequency of AAA (Lys), GAA (Glu), GAT (Asp) and AAT (Asn) codons, as compared to MAPKi-downregulated mRNAs with a non-affected TDD (Fig. 2e, lower panel).

Collectively these observations suggested that MAPKi induced the expression level downregulation of compositionally biased mRNAs by increasing their co-translational degradation (i.e., through TDD).

### Compositional biases of MAPKi-regulated ribosomal peaks

To analyse the dynamics of ribosomes on mRNAs in response to MAPKi, we performed ribosome profiling experiments and then computed mRNA ribosomal peaks (Fig. 3a and Supplementary Fig. 3a). mRNA regions in which the local density of ribosomes was higher in control cells than in treated cells are referred to as CC peaks, and mRNA regions in which the local density of ribosomes was higher in treated cells than in control cells, are referred to as TC peaks. In total, 1281 CC peaks were detected in 984 mRNAs in control cells, and 1974 TC peaks were detected in 1509 mRNAs in treated cells (Fig. 3a and Supplementary Table 1).

**Figure 3.**
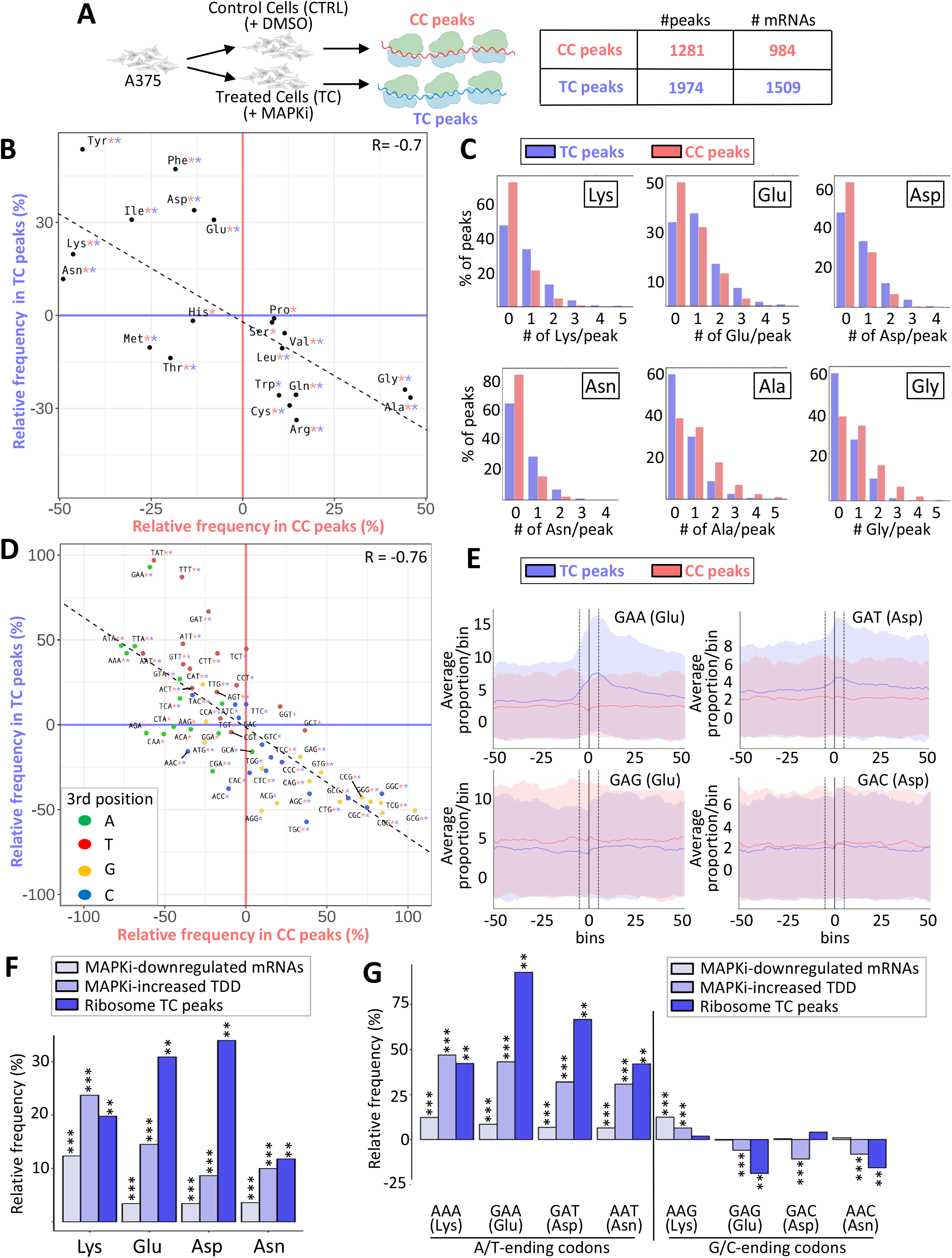
**a** A375 cells were cultured for 24h in the absence (CTRL) or in the presence of MAPKi (TC) before performing ribosome profiling and identifying ribosomal peaks in control cells compared to treated cells (CC peaks) or in treated cells compared to control cells (TC peaks). **b** Amino acid relative frequency (%) in peptides encoded by ribosome peaks in control (CC peaks) or MAPKi-treated (TC peaks) cells. The x-axis and y-axis correspond to the relative frequency (%) of each amino acid computed from CC peaks or TC peaks, respectively compared to random control peaks. * in red or in blue means that the frequency of an amino acid is statistically different (one-tailed randomization test FDR ≤ 0.05) when comparing CC peaks or TC peaks, respectively to control peaks. **c** Percentage of TC peaks (blue) and CC peaks (red) that contain different numbers (#) of Lys, Glu, Asp, Asn, Ala or Gly residues. **d** Codon relative frequency (%) in mRNA regions with a ribosome peak in control (CC peaks) or MAPKi-treated (TC peaks) cells. The x-axis or y-axis correspond to the relative frequency (%) of each codon computed from CC peaks or TC peaks, respectively compared to random control peaks. * in red or in blue means that the frequency of a codon is statistically different (one-tailed randomization test FDR ≤ 0.05) when comparing CC peaks or TC peaks, respectively to control peaks. Green, red, orange, and blue dots represent A-, T-, G-, and C-ending codons, respectively. **e** Frequencies of codons within and around ribosome peaks. The average frequencies of codons at bin 0 was computed in ribosome protected mRNA regions. The same procedure was applied for other bins (windows of 10 codons) starting from the central coordinate of each peak. The red curve corresponds to the values computed from CC peaks and the red shadow reflects the standard deviation of the values. The blue curve corresponds to the values computed from TC peaks and the blue shadow reflects the standard deviation of the values. **f** Amino acid relative frequency (%) in proteins encoded by MAPKi-downregulated mRNAs, mRNAs whose TDD increased in response to MAPKi, and in peptides encoded by regions with ribosomal peaks induced by MAPKi (TC peaks) as described in Fig. 1B, 2B, and 3B respectively. ***FDR ≤ 0.05 (beta regression analysis followed by a Tukey’s test (pairwise comparison) and ** FDR ≤ 0.05 (one tailed randomization test). **g** Codon relative frequency (%) in MAPKi-downregulated mRNAs, mRNAs whose TDD increased in response to MAPKi, and in mRNAs regions with ribosomal peaks induced by MAPKi (TC peaks) as described in Fig. 1D, 2C, and 3D respectively. ***FDR ≤ 0.05 (beta regression analysis followed by a Tukey’s test (pairwise comparison) and ** FDR ≤ 0.05 (one-tailed randomization test).

We next analysed the amino acid- and codon-composition of CC peaks and TC peaks. Some amino acids were enriched in TC peaks but reduced in CC peaks and conversely some amino acids were enriched in CC peaks but reduced in TC peaks (Fig. 3b and Supplementary Fig. 3B). For example, TC peaks contained more frequently at least one Lys, Asp, Glu and/or Asn residue than CC peaks, and the latter contained more frequently at least one Ala and/or Gly residue (Fig. 3c). Furthermore, some codons were more frequent in TC peaks than in CC peaks, while other were more frequent in CC peaks (Fig. 3d and Supplementary Fig. 3b). Interestingly, most codons enriched in TC peaks were A/T-ending codons, while most codons enriched in CC peaks were G/C-ending codons (Fig. 3d). This observation suggested that ribosomes could spend more time on A/T-ending codons in MAPKi-treated cells compared to control cells in agreement with a recent report showing that the MAPK pathway modulates codon optimality of A/T-ending codons^8^. In addition, a higher enrichment of TC peaks in the A/T-ending codons, like GAA (Glu), GAT (Asp), AAT (Asn), and AAA (Lys) was observed in contrast to the corresponding G/C-ending codons (Fig. 3e and Supplementary Fig. 3c). Finally, in agreement with a relationship between local density of ribosomes and translation-dependant mRNA decay, we observed that the TDD index of most MAPKi-downregulated mRNAs containing TC peaks increased in response to MAPKi (Supplementary Fig. 3d).

To summarize, amino acids such as Lys, Glu, Asp and Asn were enriched in i) MAPKi-downregulated mRNAs, ii) mRNAs whose TDD increased in response to MAPKi, and iii) MAPKi-induced ribosomal peaks (Fig. 3f). Furthermore, only the A/T-ending codons (AAA, GAA, GAT, AAT) corresponding to these amino acids were enriched at the expense of the corresponding G/C-ending codons, with the exception of the AAG (Lys) codon (Fig. 3g). These data support a model in which MAPKi treatment affects the dynamics of ribosomes when going through A/T-ending codons corresponding to certain amino acids (e.g., Glu and Asp), which could trigger a selective-mRNA TDD-dependant degradation.

### Amino acid bioavailability and codon-dependent selective effects

As the decrease in the intracellular concentration of certain amino acids can induce ribosome pauses and as the MAPK pathway in melanoma cells can affect amino acid metabolism (see Introduction), we next measured the intracellular concentration of Lys, Glu, Asp, Asn, Gln and Arg in the absence or presence of MAPKi. While the intracellular concentration of Lys, Gln and Arg was not affected, the intracellular concentrations of Glu, Asp and Asn decreased in response to MAPKi (Fig. 4a). This result raised the possibility that MAPKi could have a selective effect on mRNAs whose translation requires a relatively large amount of specific amino acids like Glu and Asp.

**Figure 4.**
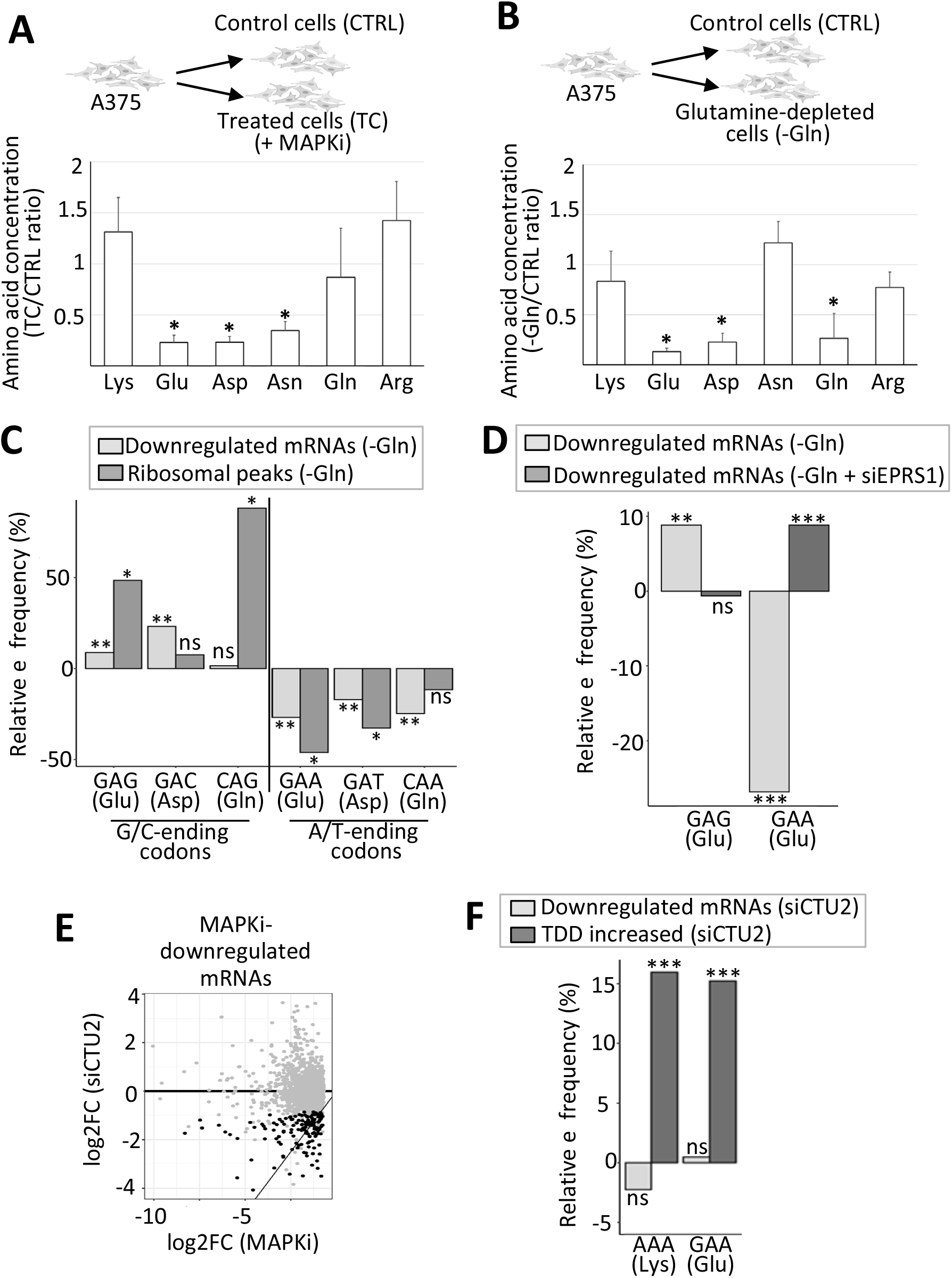
**a** Ratio of the intracellular concentration of Lys, Glu, Asp, Asn, Gln, and Arg in cells exposed 24h to MAPKi as compared to control cells. * P≤ 0.05 (two-tailed paired t-test, n=4). **b** Ratio of the intracellular concentration of Lys, Glu, Asp, Asn, Gln, and Arg in cells grown for 24h in the absence of Gln compared to control cells. * P≤0.05 (two-tailed paired t-test, n=3). **c** Relative frequency of G/C-ending codons (GAG, GAC, and CAG) and A/T-ending codons (GAA, GAT, and CAA) corresponding to Glu, Asp, and Gln in mRNAs whose expression level was downregulated in glutamine-depleted cells (-Gln) compared to all other expressed mRNAs and in ribosomal peaks induced by Gln-depletion compared to random control peaks. ** corresponds to a Tukey’s test (pairwise comparison, FDR ≤ 0.01) and *corresponds to a one-tailed randomization test (FDR≤ 0.01). **d** Relative frequency of the GAG and GAA codons corresponding to Glu in mRNAs whose expression level was downregulated in Gln-depleted cells (-Gln) and in mRNAs whose expression level was downregulated in Gln-depleted cells transfected with an siRNA targeting EPRS1 (-Gln + siEPRS1). Relative frequencies were computed against all other expressed mRNAs. Beta regression analysis followed by a Tukey’s test (pairwise comparison) FDR ≤ 0.01 (**) or < 0.001 (***). **e** The log2 fold change of the expression level of each MAPKi-downregulated mRNAs (x-axis) was plotted against the log2 fold change of their expression level in siCTU2-transfected cells compared to control cells (y-axis). Black dots represent mRNAs whose expression level was significantly (DESeq2 adjusted p-values≤ 0.05, n=3) decreased by at least 50% (while still having an average normalized expression level greater than 10) when comparing siCTU2-transfected cells to control cells. **f** Relative frequency of the A/T-ending codons (AAA and GAA) corresponding to Lys and Glu in mRNAs whose expression level was downregulated and in mRNAs whose TDD increased in siCTU2-transfected cells. *** beta regression analysis followed by a Tukey’s test (pairwise comparison, FDR < 0.001).

To challenge this possibility, we cultured melanoma cells in the absence of Asp and/or Glu. However, we did not observe any significant effect on neither Asp- and Glu-intracellular concentration, nor on cell viability (Supplementary Fig. 4a and 4b). One possible explanation is that Glu and Asp are produced from Gln provided by the growth medium. Supporting this possibility, Gln depletion from the growth medium decreased the intracellular concentration of Gln as well as Glu and Asp (Fig. 4b).

Since Gln deprivation somehow mimics the decrease of Glu and Asp intracellular concentration as observed in response to MAPKi (compared Fig. 4a, b), we analyzed the effect of Gln deprivation on gene expression and ribosome profile (Supplementary Table 1). The decrease in the intracellular concentration of Gln, Glu and Asp that was induced by Gln deprivation was associated with an enrichment of some codons corresponding to these amino acids in downregulated mRNAs and in ribosome peaks induced by Gln depletion (Fig. 4c). However, in contrast to what we observed in MAPKi-treated cells, the decrease in the intracellular concentration of Glu and Asp that was induced by Gln depletion was associated with an enrichment of the G/C-ending codons (i.e., GAG and GAC) and not the A/T-ending codons (i.e., GAA and GAT) (Fig. 4c). These results suggested that the decrease in amino acid bioavailability is not sufficient to explain codon-selective effects in agreement with previous reports (see Introduction).

Since aminoacyl-tRNA synthetases contribute to codon-selective effects, we inspected our RNA-seq datasets and we found that the expression level of several aminoacyl-tRNA synthetases varied in response to MAPKi treatment or in response to Gln depletion. Among these, we focused on the EPRS1 aminoacyl-tRNA synthetase, which loads Glu onto the corresponding tRNAs, because the expression level of EPRS1 was repressed in response to MAPKi but increased in response to Gln depletion, as validated by RT-qPCR (Supplementary Fig. 4c). To test the potential role of EPRS1 on codon-selective effects, A375 cells were cultured in the absence or presence of Gln and in the absence or presence of EPRS1 (Supplementary Table 1). EPRS1 depletion abolished the codon-selective effect of Gln depletion with respect to the enrichment of the GAG codon to the advantage of the GAA codon suggesting that EPRS1 could at least in part participate to the codon-selective effect observed after Gln depletion (Fig. 4d).

We also observed in our datasets that some enzymes that can modulate codon-selective effects by biochemically modifying tRNAs (see Introduction) were differentially expressed in MAPKi-treated cells as compared to control cells. Among these, CTU2 caught our attention for three reasons: i) CTU2 expression was decreased in MAPKi-treated cells (Supplementary Fig. 4d); ii) an important role of CTU2 in melanoma cells has already been reported^18^; and iii) CTU2, which modifies uracil on position 34 of some tRNAs, modulates the interactions between anti-codons and some A-ending codons, in particular the AAA (Lys) and GAA (Glu) codons^18^. Since we observed an enrichment of these codons in MAPKi-downregulated mRNAs, TDD-induced mRNAs, and MAPKi-induced ribosomal peaks (Fig. 3g), we tested whether the CTU2 depletion could mimic the selective MAPKi-effect. Supporting such a possibility, we first observed that CTU2 depletion resulted in the downregulation of a subset of mRNAs that were also downregulated in response to MAPKi (Fig. 4e and Supplementary Table 1). In addition, the AAA (Lys) and GAA (Glu) codons were enriched in mRNAs whose TDD was increased in CTU2 depleted cells, as expected^18^ and as observed in MAPKi treated cells (compare Fig. 4f to Fig. 3g).

In sum, our observations support a model in which the decreases in Glu and Asp bioavailability in MAPKi-treated cells resulted in an increase in the local density of ribosomes going through mRNA regions that require these amino acids to be translated (Fig. 3), which could result in translation-dependent mRNA degradation (Fig. 2), and consequently in the decrease of the expression levels of mRNAs encoding for compositionally biased proteins (Fig. 1). However, selective effects of codons ─ notably A/T-ending codons corresponding to Asp and Glu ─ probably depend on several parameters, such as the expression of aminoacyl transferases or tRNA-modifying enzymes (Fig. 4, see Discussion).

### Protein amino acid composition biases and cellular functions

Since MAPKi treatment triggers TDD of a subset of mRNAs (Fig. 2) and since TDD is likely to be dynamic and reversible, we next wondered whether the MAPKi-dependent mRNA downregulation persists after MAPKi removal, i.e., in the so-called persister cell population. To address this question, we performed RNA-seq on persister cells (Fig. 5a and Supplementary Table 1). A large number of TDD-downregulated mRNAs that we identified in cells exposed to MAPKi (Fig. 2) was still significantly downregulated in persister cells (Fig. 5a). In addition, the MAPKi-dependent decrease in Glu-, Asp- and Asn-intracellular concentration observed in MAPKi-exposed cells was also observed in persister cells (compare Figs. 4a and 5b). Furthermore, proteins encoded by MAPKi-downregulated mRNAs that were still downregulated in persister cells were enriched in Lys, Glu, Asp and Asn (Fig. 5c, left panel) and these mRNAs were enriched in the AAA (Lys), GAA (Glu), GAT (Asp) and AAT (Asn) codons (Fig. 5c, right panel). In sum, at least some composition biases observed in mRNAs that were downregulated in the presence of MAPKi were also observed in persister cells after MAPKi removal.

**Figure 5.**
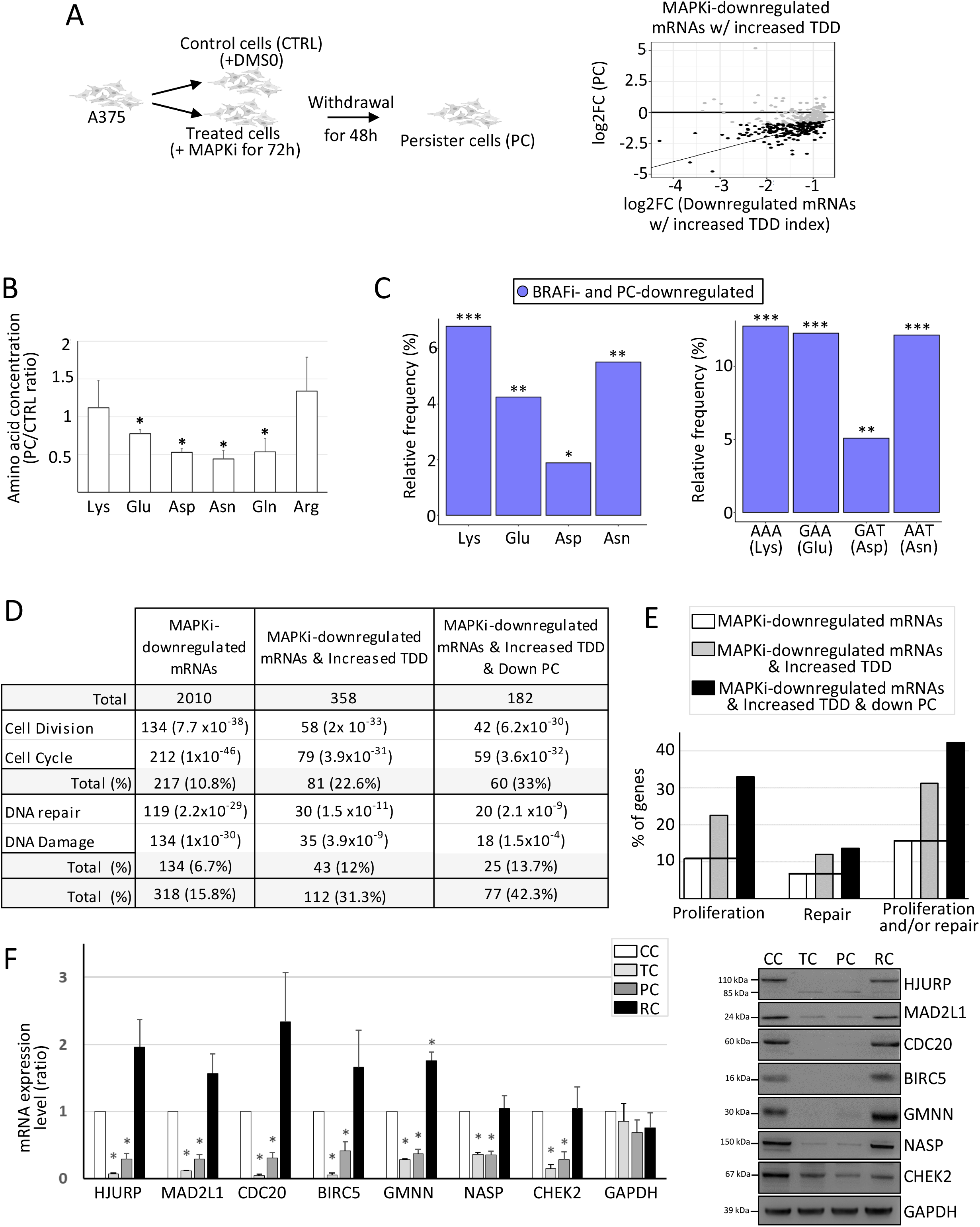
**a** A375 cells were cultured for 72h in the absence of MAPKi (CTRL) or in the presence of MAPKi (TC). Some treated cells were next grown for a supplementary 48h in the absence of MAPKi (persister cells, PC). The log2 fold change of each MAPKi-downregulated and TDD-induced mRNAs (x-axis) was plotted against the log2 fold change of their expression level when comparing persister cells (PC) to control cells (y-axis). Black dots represent mRNAs whose expression level was significantly (DESeq2 adjusted p-values ≤ 0.05, n=3) decreased by at least 50% (while still having an average normalized expression level greater than 10) when comparing persister cells to control cells. **b** Ratio of the intracellular concentration of Lys, Glu, Asp, Asn, Gln, and Arg in persister cells (PC) compared to control cells (CTRL). *P< 0.05 (two-tailed paired t-test, n=3). **c** Relative frequency (%) of Lys, Glu, Asp and Asn (left panel) computed from proteins encoded by mRNAs downregulated in both MAPKi treatment and persister cells when compared to proteins encoded by all other expressed mRNAs. Relative Frequency (%) of the AAA, GAA, GAT, and AAT codons (right panel) of mRNAs downregulated in both MAPKi treatment and persister cells when compared to all other expressed mRNAs. ***FDR<0.001, **FDR≤0.01, *FDR≤0.05 (beta regression analysis followed by a Tukey’s test (pairwise comparison)). **d** Number and functional term analysis of genes whose i) mRNAs were downregulated in response to MAPKi, ii) mRNAs were downregulated in response to MAPKi and whose TDD was increased, and iii) mRNAs were downregulated in response to MAPKi and whose TDD was increased and that were downregulated in persister cells. **e** Percentage of genes that are associated with the proliferation and/or DNA repair cellular functions. The black line indicates the % obtained from the MAPKi-downregulated mRNA population. **f** RT-qPCR and western blot analysis in control cells (CC), cells treated for three days with MAPKi (TC), cells treated for three days with MAPKi before being cultured in drug-free medium for two days (PC) or for 9 days (RC). * P< 0.05 (two-tailed paired t-test, n=3).

We next questioned the biological functions of the proteins encoded by the different mRNA populations. Some terms, like cell division, cell cycle, DNA repair and DNA damage, were enriched among the biological functions associated with MAPKi-downregulated gene products (Fig. 5d). Very interestingly, we noticed that the proportion of gene products associated with the proliferation and DNA repair biological functions increased among MAPKi-downregulated gene products whose TDD increased in response to MAPKi and that were still downregulated in persister cells (Fig. 5d, e). For example, ∼11% of MAPKi-downregulated gene products were involved in cell proliferation and this proportion reached ∼33% in MAPKi-downregulated gene products with an increased TDD and that were still downregulated in persister cells (Fig. 5d, e). Consequently, >40% of the MAPKi-downregulated gene products with an increased TDD that were still downregulated in persister cells were involved in DNA metabolism (i.e., proliferation and/or DNA repair; Fig.5d, e). This result was validated by RT-qPCR and Western blot analysis since the expression level of genes involved in cell proliferation was higher in control cells (CC) compared to MAPKi-treated cells (TC) and persister cells (PC) (Fig. 5f). Of note, the expression level of pro-proliferative genes increased after nine days of MAPKi-removal (RC, Fig. 5f), a time at which persister cells gave rise to a proliferative cell population similar to the initial one.

Because of these observations, we analyzed the composition biases of mRNAs encoding proteins involved in proliferation and DNA repair. Proteins involved in proliferation and DNA repair were enriched in a subset of amino acids, including Lys, Glu, Asp and Asn, as compared to the human proteome (Fig. 6a, b). In addition, mRNAs encoding proteins involved in proliferation and DNA repair were enriched in the AAA (Lys), GAA (Glu), GAT (Asp), and AAT (Asn) codons (Fig. 6b, left panel). Worth noting, the AAA (Lys), GAA (Glu), GAT (Asp), and AAT (Asn) codons were more enriched in MAPKi-downregulated gene products involved in proliferation and/or DNA repair when compared to the other MAPKi-downregulated gene products (Fig. 6c).

**Figure 6.**
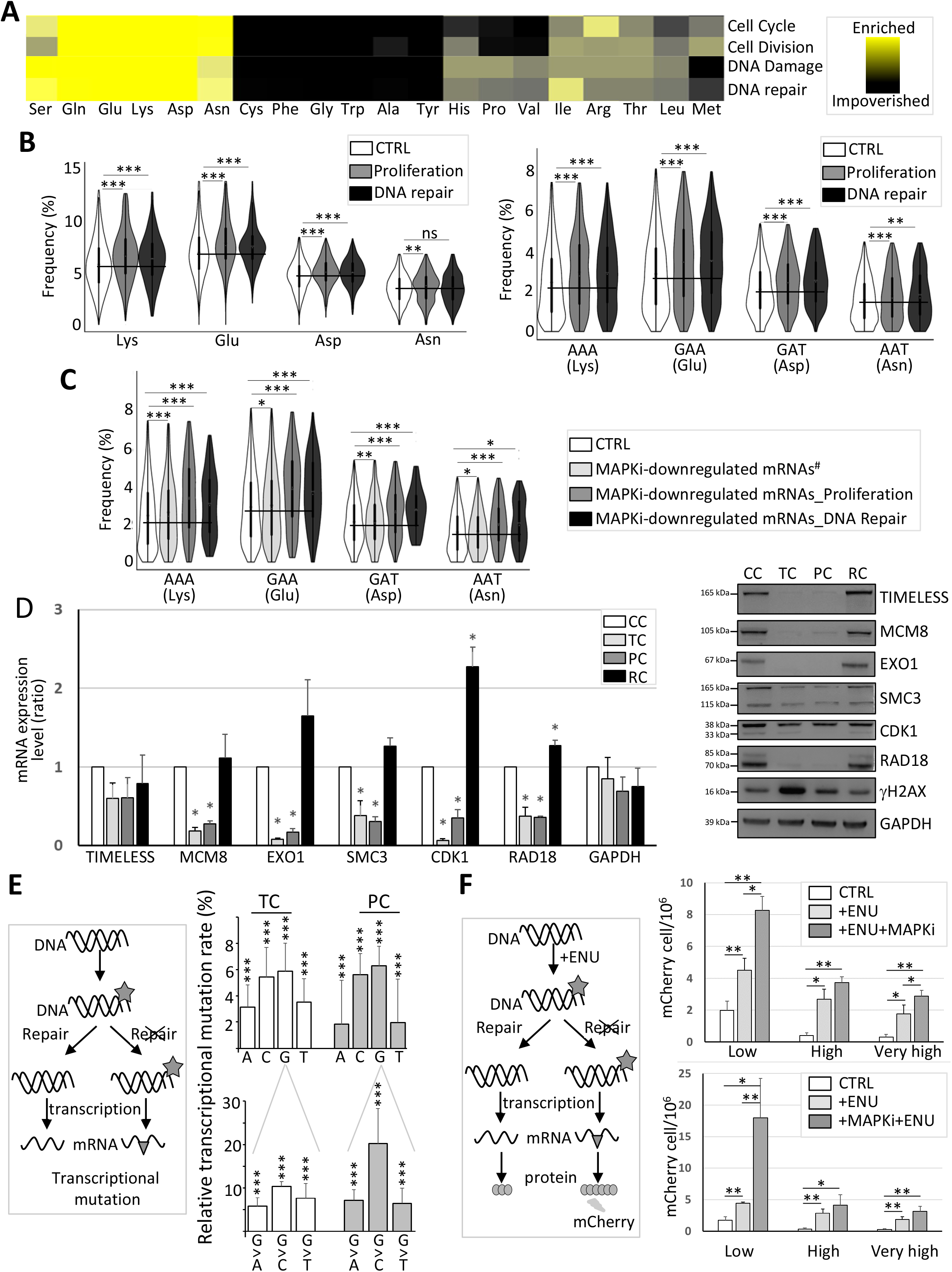
**a** Heat map representing the amino acid relative frequency in proteins involved in different cellular functions as indicated when compared to amino acid average frequency in the human proteome. **b** Frequency of amino acids (left panel) and codons (right panel) computed from control genes (CTRL) or genes involved in proliferation or in DNA repair. The black line indicates the CTRL mean value. ***FDR<0.001 and **FDR≤0.01 (beta regression analysis followed by a Tukey’s test (pairwise comparison). NS: not statistically significant. **c** Frequency of codons computed from control (CTRL) mRNAs, MAPKi-downregulated mRNAs encoding proteins not involved in proliferation or replication (MAPKi-downregulated mRNAs#), MAPKi-downregulated mRNAs encoding proteins involved in proliferation (MAPKi-downregulated mRNA_Proliferation), or MAPKi-downregulated mRNAs encoding proteins involved in DNA repair (MAPKi-downregulated mRNA_DNA Repair). The black line indicates the CTRL mean value. ***FDR <0.001, **FDR≤0.01, *FDR≤0.05 (beta regression analysis followed by a Tukey’s test (pairwise comparison)). NS: not statistically significant. **d** RT-qPCR and western blot analysis in control cells (CC), cells treated for 3 days with MAPKi (TC), cells treated for 3 days with MAPKi before to be grown in drug-free medium for 2 days (PC) or for 9 days (RC). *P<0.05 (two-tailed paired t-test, n=3). **e** The schematic representation on the left describes how DNA damages (grey asterisk, e.g. DNA chemical modifications) can lead, in the absence of DNA repair, to transcriptional mutations (grey triangle) owing to nucleotide mispairing. Transcriptional mutations were quantified by comparing at the nucleotide level the transcriptome of control cells to the transcriptome of MAPKi-treated cells (TC) or persister cells (PC). The top panel represents the relative % of A, C, G, and T nucleotides that are more frequently mutated to another nucleotide in MAPKi-treated cells (TC) or persister cells (PCs) compared to control cells. The bottom panel represents the relative % of G nucleotides that are more frequently mutated to another nucleotide in MAPKi-treated cells (TC) or in persister cells (PC) when compared to control cells. *** Logistic regression analysis FDR < 0.001 (n=3). **f** The schematic representation on the left describes how DNA damages (grey asterisk, e.g. DNA chemical modifications induced by ENU) can lead, in the absence of DNA repair, to nucleotide mispairing that transforms a stop codon (preventing mCherry synthesis) into a tryptophan codon (allowing mCherry synthesis). The top panel represents the quantification of the number of cells expressing mCherry either under control conditions (CTRL), after exposure to ENU (+ENU), or after exposure to ENU in the presence of MAPKi (+ENU+MAPKi) (n=4). The bottom panel represents the quantification of the number of cells expressing mCherry either under control conditions (CTRL), after exposure to ENU (+ENU), or after exposure to MAPKi for 72h before being exposed to ENU (+MAPKi+ENU) (n=5). Low, high, and very high correspond to different filters used in cytometric analysis to detect positive cells (i.e. cells expressing mCherry). *P<0.05 and **P<0.01 (one-tailed paired t-test).

Since the expression level of mRNAs encoding proteins involved in DNA repair decreased in MAPKi-treated cells and in persister cells, as validated by RT-qPCR and Western blot analysis, while γ-H2AX ─ a marker of DNA damage ─ was significantly increased in MAPKi-exposed cells (Fig. 6d), one hypothesis is that MAPKi-exposed cells may have a higher probability of accumulating DNA damage that could increase the probability of genetic mutations to appear in descendant cells. However, measuring the genetic mutational rate of MAPKi-exposed cells is challenging, as genetic mutations can only be quantified after several rounds of replication cycles, while MAPKi represses cell proliferation. In order to circumvent this difficulty, we used two complementary approaches.

First, we decided to look for nucleotide variations within mRNAs. Indeed, genetic mutations are notably the consequence of nucleotide mismatches during replication that themselves are the consequence of nucleotide chemical modifications (i.e., DNA damage such as nucleotide oxidation). In this setting, nucleotide chemical modifications can also lead to nucleotide mismatches during transcription, giving rise to the so-called transcriptional mutations^52–58^. In other words, unrepaired DNA damage ─ such as nucleotide chemical modifications ─ can lead to transcriptional mutations in neo-synthetized mRNAs. By comparing the transcriptome of MAPKi-exposed and persister cells to control cells at the nucleotide level, we observed that the relative frequency of transcriptional mutations affecting each of the four nucleotides increased both in MAPKi-treated cells (TC) and in persister cells (PC) (Fig. 6e, upper panel). For example, there was between 5% to 20% more transcriptional mutations changing a G nucleotide into A, C or T in MAPKi-treated cells (TC) or persister cells (PC) when compared to control cells (Fig. 6e, lower panel).

The second approach we used was to generate a cellular clone expressing a reporter gene that contains a TGA stop codon preventing the synthesis of the mCherry protein, that can only be expressed if nucleotide biochemical modifications ─ induced for example by mutagenic agents, such as ENU ─ result in nucleotide mismatches that change the TGA stop codon into the TGG codon coding for tryptophan^59^. As expected, the number of mCherry-positive cells was increased after ENU treatment regardless of the filters used in cytometry to count positive cells (Fig. 6f, upper panel). Importantly, the ENU effect was increased when cells were simultaneously exposed to MAPKi (Fig. 6f, upper panel) or when cells were first exposed to MAPKi before to be exposed to the ENU (Fig. 6f, lower panel). Collectively, our results point to a link between the downregulation of DNA repair genes in MAPKi-exposed cells and a higher rate of nucleotide mismatches.

## Discussion

The compositional biases of MAPKi-downregulated gene products that we observed (Fig. 1) could be explained at least in two ways. First, from a gene centric point of view, the MAPK pathway could have evolved to repress the transcription of genes that are involved in DNA metabolism such as DNA replication. As gene products involved in DNA metabolism bear compositional biases (Fig. 6a, b; see below), then the MAPKi-downregulated gene products would bear these function-related compositional biases. The second possible explanation, which we term a metabolic centric point of view, is that the MAPKi-dependent decrease of the bioavailability of some amino acids (e.g., Asp and Glu) results in the translation-dependent expression level decrease of Asp- and Glu-enriched gene products. Since proteins involved in DNA metabolism are enriched in Asp and Glu, then MAPKi treatment induces the expression level decrease of gene products involved in DNA metabolism in a metabolism-depending manner. As discussed below, our results and those from previous publications support this metabolic centric point of view without excluding the well-established effects of MAPKi on the transcriptional activity of genes involved in cell proliferation.

The BRAF^V600E^ mutation-dependent hyper-activation of the MAPK pathway in melanoma cells triggers not only cell proliferation but also the cellular addiction to some non-essential amino acids, such as Gln. Indeed, in mutated melanoma cells, the carbon skeleton of Gln fuels both non-oxidative energetic metabolism and amino acid and nucleotide biosynthetic pathways on which growing cell depends^35, 45, 51, 60–63^. Accordingly, growth medium depletion of Gln induced Glu- and Asp-intracellular concentration decrease, while Glu- and/or Asp-depletion did not result in their intracellular concentration decrease (Fig. 4b and Supplementary Fig. 4b), probably because Glu and Asp can be generated from Gln from the growth medium^45, 63^. Importantly, the use of Gln, Glu, and Asp in the oxidative phosphorylation (OXPHOS) pathway, which produces energy from the complete degradation of their carbon skeleton, is reactivated by MAPKi^46–51, 60^. This may explain the observed Glu- and Asp-intracellular concentration decrease in MAPKi-exposed cells (Figs. 4a and 5b). This together with the observed enrichment of these amino acids in i) MAPKi-downregulated mRNAs (Fig. 1b), ii) MAPKi-induced TDD mRNAs (Fig. 2b), and iii) MAPKi-induced ribosomal peaks (Fig. 3b) support a model where MAPKi-dependent effects on the cellular metabolism impacts the translation-dependent mRNA expression level through the bioavailability of amino acids.

While the precise codon composition bias of MAPKi-downregulated gene products is likely depending on several parameters (Fig.4 and see Introduction), the observed amino acid composition bias of MAPKi-downregulated gene products is particularly interesting as it may explain how cells “coordinate” their metabolic activity and their phenotype. Indeed, we observed that gene products involved in DNA metabolism (e.g., DNA replication) are enriched in charged amino acids such as Glu and Asp, which can be explained in different ways. First, proteins involved in DNA metabolism are hydrophilic proteins, which rely on protein enrichment in hydrophilic amino acids such as charged amino acids (Supplementary Fig. 1a). Second, proteins involved in DNA metabolism contain positively- and negatively-charged amino acids that play an important role in protein-DNA interaction^64–66^. Finally, proteins involved in DNA metabolism contain negatively-charged amino acids that interact with ions (e.g., Mg^2+^) on which depends their enzymatic activities^67^. Since charged-amino acids like Glu and Asp are at the crossroad between energetic- and gene product synthesis-pathways, their degradation through the OXPHOS pathway would decrease their bioavailability, with consequences on the translation of gene products involved in cell proliferation that require these amino acids. Therefore, cells that survive MAPKi treatment would be the cells that consume amino acids through the OXPHOS pathway, which simultaneously reduced their needs in terms of amino acids and nucleotides because of the metabolic-dependent lower biosynthesis of proliferation gene products. Accordingly, persister cells (i.e., non-genetically modified cells that survive anti-MAPK therapy) i) have a lower intracellular concentration of Glu and Asp as compared to the initial cell population (Fig. 5b), in agreement with the OXPHOS pathway re-activation reported in these cells, and ii) express a low level of gene products involved in DNA metabolism (Fig. 5d–f), in agreement with their reported slow growing rate^46–48, 68–73^.

While favoring survival versus proliferation, the MAPKi-dependent decrease in the intracellular concentration of Glu and Asp could have a secondary effect because of the expression level decrease of gene products involved in DNA repair (Figs. 5d, e and 6d), that share the same composition biases with genes involved in cell proliferation (Fig. 6a–c). The expression level decrease of gene products involved in DNA repair may seem to be of no consequence in non-dividing cells since genetic mutations can only occur during replication. However, the fact that unrepaired DNA damage can induce base-pairing mismatches during transcription ─ leading to transcriptional mutations^52–58^ ─ could explain the observed larger number of nucleotides variations in the transcriptome of cells exposed to MAPKi compared to control cells (Fig. 6e). Interestingly, it has been proposed that transcriptional mutations could be a “pre-selection step” toward the emergence of genetically-modified and –adapted cells^52–58^. Indeed, if unrepaired DNA damage leads to the synthesis of mutated gene products that contribute to the survival of a cell, this cell may have a higher probability of generating genetically-modified and – adapted descendant cells because the same unrepaired DNA damage could trigger a genetic mutation. Although this model is speculative with respect to the data we provided, persister cells have been proposed to be a reservoir of genetically modified and therapy-resistant cells^69^.

In conclusion, we propose that MAPKi-induced metabolic changes result in the bioavailability decrease of amino acids such as Glu and Asp, which contributes to the expression level decrease of proteins enriched in these amino acids, including proteins involved in proliferation and DNA repair (Fig. 7). The coupling between metabolism and gene expression could, as a side effect, results in the accumulation of DNA damage ─ owing to the expression level decrease of DNA repair enzymes ─ leading first in transcriptional mutations and then in genetic mutations, increasing therefore the probability of genetically-mutated and –adapted clones to emerge in response to MAPKi.

**Figure 7.**
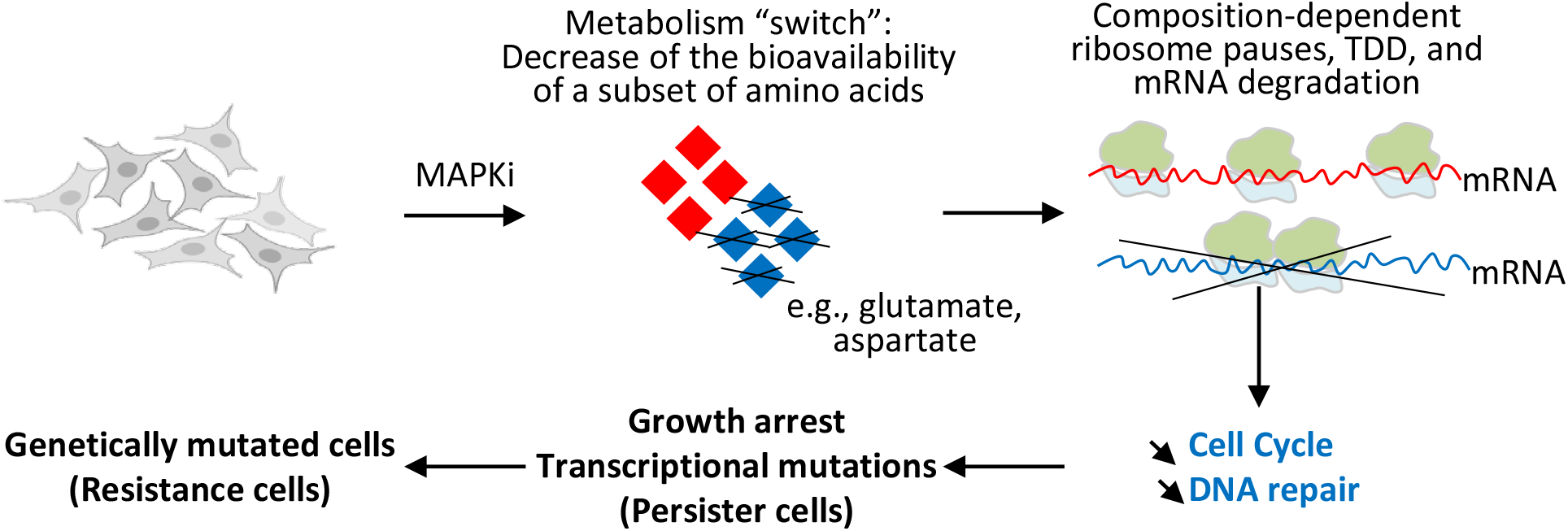
By switching the cell metabolism and decreasing the bioavailability of certain amino acids such as glutamate and aspartate, MAPKi could trigger ribosome pause sites on some mRNA regions enriched for codons corresponding to glutamate and aspartate, which in turn could trigger the selective degradation of a subset of mRNAs according to their compositional biases in certain codons and corresponding amino acids. Since the biological functions of proteins depend on their composition in certain amino acids, the selective degradation of mRNAs according to their compositional bias would affect a selective set of functions such as proliferation and DNA repair. Since, the downregulation of compositionally-biased gene products persists in cells after MAPKi withdrawal, the selective degradation of compositionally-biased mRNAs could simultaneously contribute to the appearance of slow-proliferative cells that would have a higher probability to generate mutated daughter cells.

## Methods

### Cell culture and persister cells

The human melanoma A375 cell line (ATCC) was cultured at 37 °C and 5% CO^2^ in Dulbecco’s Modified Eagle’s medium (DMEM; Gibco) supplemented with 10% FBS, 2 mM glutamine and penicillin– streptomycin. Cells were split at 80% confluence, three times a week. MAPKi-treated cells were cultured in a medium containing 1 µM vemurafenib/cobimetinib (with 500 nM vemurafenib and 500 nM cobimetinib) (Euromedex) for 24 h. Glutamine-depleted cells were cultured in a glutamine-free medium for 24 h. Persister cells were obtained after being cultured for 72h in a medium containing 1 µM vemurafenib/cobimetinib in three experimental batches (A–C). Batch A was harvested after three days and batches B and C were cultured in drug-free DMEM for two and nine additional days.

### siRNA transfections, cell harvesting, RNA extraction and qRT-PCR

Two different siRNAs (Merck, Supplementary Table 2) that target constitutive exons with minimum predicted off-targets were designed and pooled together. Cells were reverse-transfected with lipofectamine RNAiMAX (ThermoFisher), following the manufacturer’s instructions. After transfection, cells were washed twice with ice-cold PBS, scraped with 1 ml of PBS and pelleted by centrifugation (500×*g* for 1 min at 4 °C). Cells were then suspended in 1 ml lysis buffer (10 mM Tris-HCL pH 7.5, 5 mM MgCl_2_, 100 mM KCl, 1% Triton X-100) and incubated on ice for 10 min. Cellular lysates were centrifuged (1000×*g* for 10 min, 4 °C), and the cytoplasmic supernatants were used for the subsequent experiments. RNA was extracted using TRI Reagent (Sigma), following the manufacturer’s instructions. For RT-qPCR, 1 µg of extracted RNA was retro-transcribed using the Maxima First Strand cDNA Synthesis Kit (ThermoFischer), following the manufacturer’s instructions. qPCR reactions were run in triplicate on a LightCycler 480 (Roche) in 10 µl reactions. The amino acid intracellular concentration was performed by the AltaBioscience and Xell companies.

### Gene annotation

Bioinformatics analyses were performed using the GRCh38.p13 assembly and NCBI’s annotation. Only genes with at least one coding sequence (CDS) and one start and stop codon were kept. Merged genes, and CDS that overlap several exons or an ambiguous coding frame were filtered out. A total of 19,143 coding genes and 196,652 CDS were selected. In the following analyses, the mRNAs are the concatenation of the CDS of a gene and a gene was associated with only one mRNA.

### RNA-seq and QuantSeq

RNA-seq libraries were prepared and sequenced by Novogene (rRNA depletion library preparations, sequencing on Novaseq 6000 2x150). QuantSeq libraries were prepared using the QuantSeq 3’ mRNA-Seq Library Prep Kit (Lexogen). RNA (500 ng) was spiked-in with 1 µl of a 1:100 dilution of ERCC Spike-In Mix (ThermoFisher) prior to library preparation. QuantSeq libraries were then quantified, pooled and sent for sequencing at Novogene (Novaseq 6000 2x150). Only fastq files containing forward reads were used for QuantSeq analyses, while the entire pair of fastq files were used for RNA-seq analyses. Adapters were removed from raw reads and trimmed using fastp version 0.20.1^74^ with the following parameters: qualified_quality_phred 30 -l 25 --detect_adapter_for_pe) (RNA-seq) and qualified_quality_phred 20 -3 --cut_tail_window_size 10 -- adapter_sequence=AGATCGGAAGAGCACACGTCTGAACTCCAGTCA –adapter_sequence=AAAAAA (QuantSeq). Reads were mapped against the human genome GRCh38.p13 with HISAT2 version 2.2.1. Reads were counted on exons with htseq-count version 0.13.5. Unexpressed genes (genes with raw read counts < 2 across all conditions) were filtered out only for RNA-seq data. Differential expression analysis was performed with the DESeq2 package version 1.34.0 using the option lfcThreshold = 0.585 (only RNA-seq data). Differentially expressed genes with an average DESeq2 normalized expression above 10 across conditions were kept (only RNA-seq data).

### TDD experiments and analysis

TDD monitoring was performed as previously described (https://doi.org/10.1101/2020.10.16.341222). Briefly, the culture medium was removed at 24h after plating cells and replaced with fresh medium containing 1 µM vemurafenib/cobimetinib (with 500 nM of each) or DMSO. After another 18 h, cells were treated with fresh cycloheximide (100 µg/ml) or DMSO for 5 min and then treated with tryptolide (25 µM) or DMSO. This was prepared in four identical batches (A–D): batch A was harvested immediately upon +/- tryptolide treatment (T0 samples), and the batches B, C and D were harvested after 3 (T3) and 5 (T5) hours of treatment, respectively. Computation of the TDD index was performed using pre-processed reads, mapped and counted with htseq-count (see above). From the raw count tables obtained with htseq-count, the CPM was computed for each gene. Then, we searched for genes with a stable expression in MAPKi and DMSO conditions for normalization. A gene was considered as stable if i) its CPM count was greater than 0.2 in initial condition, ii) its CPM count after 3 or 5 hours of transcription inhibitor treatment was at least 10% greater than its CPM at initial condition; and iii) its CPM at initial condition was greater than 10% of its CPM after 3 or 5 hours of transcription inhibitor treatment. Only “stable genes” in all replicates were kept. Stable genes were used as normalization factors in DESeq2 package to normalize reads counts. The TDD index was next computed for an mRNA produced by a gene G using the following formula:

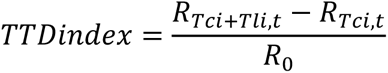

where *R*_0_ is the normalized number of *G* reads at 0h (initial condition), *R*_*Tci+Tli,t*_ is the normalized number of *G* reads after *t* hour of exposition to a transcription (Tci) and a translation inhibitor (Tli) and *R*_*Tci,t*_ is the normalized number of *G* reads after *t* hour of exposition to a Tci. The TDD index was computed, for each replicate, at T3 and T5 for cells treated with MAPKi and DMSO. The TDD index according to the condition (MAPKi or DMSO), the time (T3 or T5) and the replicate was modeled using a linear model (in R function lm) for each mRNA. With these models, a student test was computed to test if the TDD index of a mRNA increased or decreased in response to MAPKi as compared to the DMSO condition.

### Ribosome profiling and analysis

Ribosome profiling samples were prepared as described in https://doi.org/10.1101/2022.04.29.489990. Briefly, cells were harvested with PBS and lysis buffer supplemented with 100 µg/ml cycloheximide (Sigma) and 2 mM DTT. The lysate was analysed at 260 nm absorbance to estimate the total quantity of material and then treated with nucleases. For every 5 units of A260 absorbance, 6 µl of MNase (1 mg/ml; Nuclease S7, Roche) was added to the lysate, along with CaCl_2_ to a final concentration of 10 mM. Lysates were then incubated at 25 °C for 30 min, transferred to ice and then applied to 10–50% sucrose gradients containing cycloheximide (100 µg/ml). After ultra-centrifugation at 35,000 rpm for 2h and 40 min at 4 °C, gradients were fractionated using a fraction collector, and the fractions containing the digested monosome fragments (80S) were kept. Fractions were supplemented with SDS (to a final concentration of 1%) and then digested with proteinase K (Roche, final concentration of 2 µg/ml) for 45 min at 42 C. Protected RNA fragments were then purified using an acidic phenol–chloroform extraction (Fischer, BP1753I) and precipitated overnight at –20°C with 0.1× volume of sodium acetate (3 M, pH 5.2), 1× vol isopropanol, 1 µl GlycoBlue and 10 mM MgCl_2_ (to help recover smaller nucleic acids)_._ Purified RNA fragments were then 3’-end dephosphorylated using PNK and fractionated on a 10% acrylamide denaturing gel, and the smears of interest (26–32 bp) were cut from the gel and purified. Size-selected fragments were rRNA-depleted by hybridization using RNA probes^75^, successively RNAse H– and DNAse-treated and then purified again with a phenol–chloroform extraction before proceeding to cDNA library preparation following the Omniprep Library preparation protocol^76^. An adaptator sequence was ligated to RNA, which was then retrotranscribed with barcoded primers. The barcoded cDNAs were size-selected on a 10% denaturing acrylamide gel, purified and then circularized (CircLigase, Lucigen). Amplification with barcoded primers was performed with a few numbers of PCR cycles (5 to 8) and a high-fidelity polymerase (Q5, NEB). Amplified libraries were size-selected on a non-denaturing 8% acrylamide gel and purified, and their quality and concentrations were assessed using the TapeStation DNA 1000 ScreenTapes. Ribosome profiling OmniPrep libraries were sequenced by GenomEast (HiSeq 4000 1×50bp). After removing adapter sequences from raw reads using cutadapt version 2.1 with the parameters -a AGATCGGAAGAGCACACGTCTGAACTCCAGTCAC -u 13 --maximum-length=40 -- minimum-length=20 -q 28,28 for Ribo-seq data, and the parameters -a AGATCGGAAGAG -g CTCTTCCGATCT -A AGATCGGAAGAG -G CTCTTCCGATCT for RNA-seq data (raw reads, see “Total RNA-seq and QuantSeq" section). A trimming step was then performed using UrQt version 1.0.18^77^, and reads were mapped to GRCh38.p13 genes sequences using HISAT2 version 2.2.1^78^ with parameters -- rna-strandness ‘F’ --norc for Ribo-Seq data. Next, alignment files were converted, using deepTools version 3.0.2^79^, to bigWig files containing a count per million (CPM) mapped reads normalized coverage at one nucleotide resolution. A peak calling step and a statistical analysis were then performed. For a replicate *i* and a gene *G,* a normalized coverage *cNorm* was computed for a test *T* and a control *C* condition using the following formula:

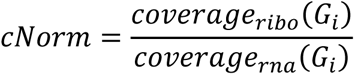

where *coverage_ribo_* was obtained from Ribo-seq data, and *coverage_rna_* from RNA-seq data. Nucleotide positions at which no coverage was detected from RNA-seq data were skipped. The difference *cDiff_i_* between normalized coverage in *T* and *C* condition was then computed for each replicate *i*. Thus, for a number *N* of replicates, a set of coverage *CDIFF* per replicates was obtained as follows:

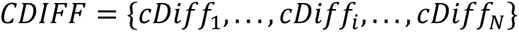

The average coverage *cMean* = {*cMean*_0_, . . . , *cMean*_*p*_, . . . , *cMean*_*L*−1_} at each CDS position *p* of a gene G of length *L was* computed between replicates, where *cMean_p_* was computed as follows:

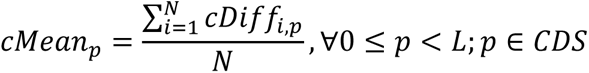

Next, the average coverage *MeanCov* and standard error *StdMean* were computed for the gene *G*, and a coverage threshold *T* was defined with *T* = *MeanCov* + (*StdMean*) × 3, where *MeanCov* and *StdMean* were computed using the formulas:

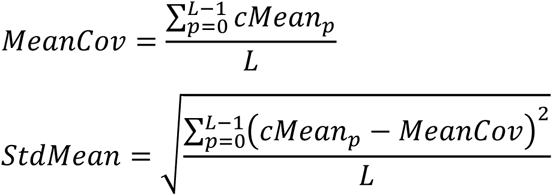

Each region at which *cMean* was above *T* was considered as a peak. Peaks inside CDS with an average coverage below 3 across RNA-seq replicates, were removed. Peaks defined by a region where the average RNA-seq coverage was below 3 were discarded. A score was given for each peak, beginning at a position *s* and ending at a position *e* in a gene. Only peaks position with a score above 3 in two replicates were kept. The score was computed using the formula below:

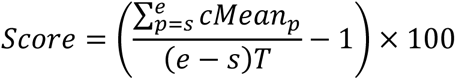

The first analysis was done by taking the MAPKi or glutamine-deprived conditions as the test condition, and the DMSO or untreated condition as the control condition, respectively. For each analysis performed with this method, another was carried out by reversing the control and test conditions.

Randomization tests were performed to test whether a set of peaks *P* had a codon compositional bias, or whether peptides encoded in peak regions had an amino acid bias. For this, 10,000 sets of control peaks *C* similar to *P* (same peak number and peak size) located in CDS were sampled. For each peak of *P* and *C*, the frequency of a given feature *X* (*i*.*e*. codon or encoded amino acid) was computed using the formula: *Freq*(*X*) = *Count* (*X*)/*s*, where *Count(X)* is the number of *X* in a peak and *s* is the total number of codons in this peak. The average frequency of *X, Mean_X_,* was then computed for *P* and the 10,000 sets of *C MEAN*_*c*_ = {*Mean*_*c*1_, . . . , *Mean*_*c*10000_}. To calculate an empirical p-value, the number of control frequencies *Mean_Ci_* upper or equal or lower or equal than the frequency *Mean_X_* was determined. The smaller number between these two was then divided by the number of control peak sets (*i.e.* 10,000). Note that the p-value cannot be lower than 1/10,000 to avoid multiple testing caveats. The p-value was then corrected using the Benjamini–Hochberg procedure. Figure 3E was generated with codons inside coding sequences (CDS) of genes producing transcripts that have at least one ribosome peak. Codons overlapping two different CDS were discarded from the analysis. The frequency of a given codon was computed in the CDS region overlapped by a peak. Starting from the central coordinate of the peak ((end - start)/2 rounded up), the frequency of codons was then computed up to 50 windows of 10 codons with a step of 1 upstream and downstream the peak.

### Compositional bias analyses

To test whether the codon content of different sets of genes was different, the frequencies of each codon in genes according to their size and set was modeled with a generalized linear model for the beta distribution with zero inflation (with R glmmTMB function of the glmmTMB package using beta_family(link = "logit") parameter). Then, a Tukey’s test (pairwise comparison) for the ‘set’ factor was done (with R emmeans and pairs functions of the emmeans package). Control sets of genes correspond to expressed genes having a mean DESEQ2 normalized expression greater than 10 and not being in other tested sets. The same procedure was applied to test whether the amino acid content of different sets of proteins was different. When several codons or amino acids are displayed in a figure, an additional Benjamini-Hotchberg correction is performed. The relative frequency of a feature X was computed as follow:

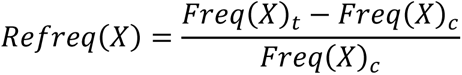

Where Freq(X)t is the average frequency of a codon or amino acid X in a test set of mRNAs or proteins and Freq(X)c is the average frequency of a codon or amino acid X in the set of mRNAs expressed in A375 cells or their encoded proteins.

### Functional enrichment analysis and heatmap

Gene ontology (GO) enrichment analysis was performed using DAVID Ontology^80^. An annotation file containing GO terms and a gene association file (that links proteins to their most specific GO terms) were downloaded from http://geneontology.org/. A homemade tool was developed to extract all proteins associated with GO:0051301 (cell division) and GO:000628 (DNA repair). Proteins associated with child terms of these GO terms were also treated as belonging to these terms. Only child terms linked to their parents with the qualifiers ‘involved_in’, ‘located_in’, ‘is_active_in’, or ‘part_of’ were considered. In addition, proteins associated with the Uniprot keywords KW-0131 (cell cycle) and KW-0227 (DNA damage) were downloaded from https://www.uniprot.org/keywords/. Only reviewed human proteins were kept. The average frequency of each amino acid in these lists of proteins was calculated using FasterDB (http://fasterdb.ens-lyon.fr/faster/home.pl). Overall, 10,000 sets of proteins were randomly sampled for each list of proteins, and the average frequency of each amino acid for each set was computed. Finally, an empirical p-value was computed for an amino acid *X* in a given protein list *P* as:

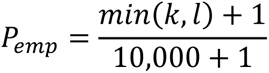

where k is the number of controls sets with an average frequency of *X* higher or equal to *P*, and *l is* the number of controls sets with an average frequency of *X* lower or equal to *P.* For each list of proteins, the p-values were corrected using the Benjamini-Hochberg procedure and then transformed using the following formula: *T* = 1 − *P*_*adj*_ × *s*, where *P*_*adj*_ is the corrected p-value and *s* =1 if *k > l;* <u>otherwise,</u> *s* = − 1.

### Transcriptional mutagenesis

Mapped reads files (see “RNA-seq and QuantSeq” section) were recovered and duplicated reads were removed using the program MarkDuplicates from picard toolkit version 2.18.11 (Picard Toolkit 2019. Broad Institute, GitHub Repository. https://broadinstitute.github.io/picard/) with the parameters VALIDATION_STRINGENCY=LENIENT REMOVE_DUPLICATES=true. Then, the number of mapped reads within each files were recovered using idxstats commands of samtools v1.11. Each bam file was sub-sampled, using samtools v1.11, to have approximately the same number of mapped reads as in the smallest bam file. The command mpileup of the program bcftools v1.16 was used to produce bcf files using the following options: -I -d 10000 -O b -a AD. The mpileup was only performed on human exonic regions. SNP and unchanging nucleotide positions were next called using the command call from bcftools and the parameters -A -V indels -m -O b. The resulting positions were filtered by depth and quality with the command filter from bcftools and the parameters -i ‘QUAL>=10 && DP>=700’ -O b. Finally, these bcf files were again filtered using a homemade Python script to keep only positions that have at least 700 nucleotides of coverage depth and an alternative allele frequency lower than 5%. A transcriptional SNP was identified by REF>ALT, where REF is the nucleotide found in the reference genome at a particular position, and ALT is the nucleotide found on mapped reads at this position with REF ≠ ALT. The number X of nucleotide positions with a coverage depth greater than 700 and containing a SNP REF>ALT was recovered. This number X was then divided by the total number of nucleotide REF with a coverage greater than 700 to obtain a proportion of sites REF with a SNP REF>ALT. Then, we tested whether the proportions of the same SNP across different conditions were different by using a logistic regression. We used the same procedure to test whether the proportion of SNP REF>* are different between conditions. A SNP REF>* corresponds to any SNP located on a given nucleotide REF with a coverage above 700 in the genome. The relative SNP frequency of a given condition compared to DMSO-treated cells was computed using the same formula as defined in “Compositional bias analyses section”.

### Mutagenesis reporter experiments

A stable clonal A375 cell line expressing GFP and a mutated and non-fluorescent version of mCherry (CherryOFF) was obtained from retro-viral particles, prepared from the pQC-CherryOFF-GFP plasmid according to Birnbaum et al^59^. pQC-CherryOFF-GFP was a gift from Fangliang Zhang (Addgene plasmid #129101 ; http://n2t.net/addgene:129101 ; RRID:Addgene_129101). Genetically-modified cells were treated for 72 h with DMSO (CTRL), 1 mM ENU (+ENU, N3385, Merck), or 1 mM ENU + 1 µM vemurafenib/cobimetinib (+ENU+MAPKi). Alternatively, cells were treated with DMSO (CTRL), 1 mM ENU (+ENU), or 1 µM vemurafenib/cobimetinib (MAPKi) for 72 h before to be cultured in a drug-free medium for 6 days before to be treated with 1 mM ENU (+MAPKi+ENU) for 72 h.

### Data availability

The raw NGS datasets from this study were deposited on Gene Expression Omnibus (GSE232604; GSE232709; GSE233122; GSE232551; GSE233120; GSE233121)

## Supplementary information

**Additional file 1: Supplementary Table 1**

**Additional file 2: Supplementary Table 2**

**Additional file 3: Supplementary Figures**

## Acknowledgements

We gratefully acknowledge the support from the PSMN (Pôle Scientifique de Modélisation Numérique) of the ENS de Lyon for the computing resources. We thank the members of the LBMC Biocomputing Hub and V.A. Raker for manuscript editing.

## Author contributions

F.A., L.G. and E.V. performed experiments; N.F. and A.L. performed bioinformatics analyses; E.P.R. and D.A. conceived the project and supervised the experimental work and the bioinformatics analyses; F.A., N.F., E.P.R and D.A. wrote the manuscript with help and comments from all co-authors; and all authors read and approved the final manuscript.

## Ethics approval and consent to participate

N/A. The cell lines were authenticated by ATCC

---

**Supplementary Figure 1.**
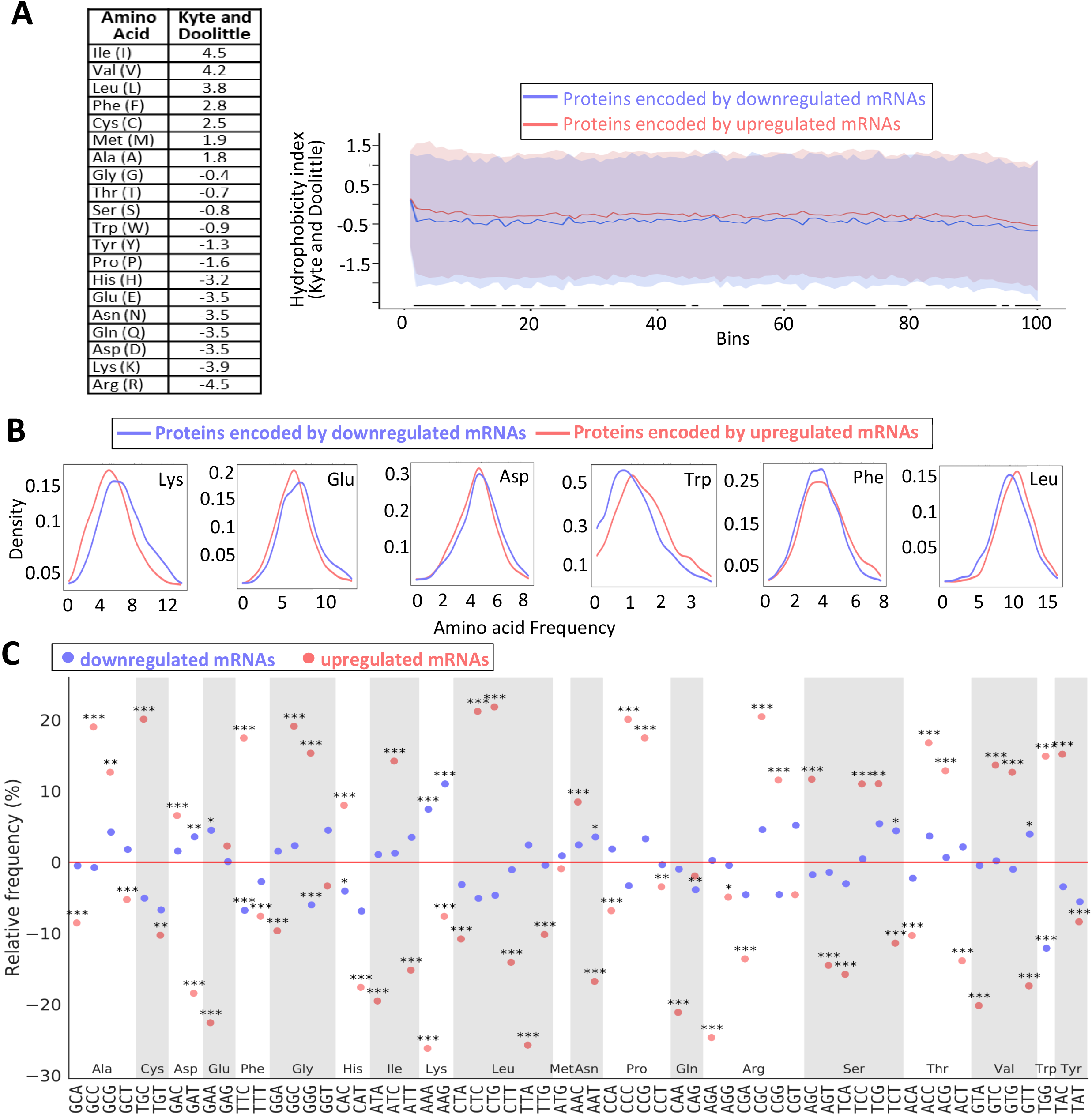
**a** Hydrophobicity index of proteins encoded by MAPKi-regulated mRNAs. On the left, hydrophobicity scale defined by Kyttle and Doolittle (“A simple method for displaying the hydropathic character of a protein” J Mol Biol. 1982 May 5;157(1):105-32.PMID: 7108955 DOI: 10.1016/0022-2836(82)90515-0). On the right, averaged-hydrophobicity index of proteins ─ divided into 100 bins ─ that were either encoded by MAPKi-upregulated mRNAs (in red) or by MAPKi-downregulated mRNAs (in blue). The red- and blue-background represent the variability (standard deviation) of the hydrophobicity index insight each of the two protein groups. The bottom black lines represent the statistical analysis (two-tailed t-test FDR ≤ 0.05) computed at each position. **b** Distribution curves of the proportion (density) of proteins encoded by MAPKi-upregulated mRNAs (red) or by MAPKi-downregulated mRNAs (blue) as a function of the frequency of Lys, Glu, Asp, Trp, Phe, or Leu. Shifting the curves to the right means that a higher proportion of mRNAs in a given mRNA population has a higher frequency of a given amino acid. **c** Relative frequency of codons in MAPKi-downregulated mRNAs and MAPKi-upregulated mRNAs when compared to the frequency of codons in A375-expressed mRNAs. ***FDR<0.001, **FDR≤0.01, *FDR≤ 0.05 (beta regression analysis followed by a Tukey’s test (pairwise comparison)).

**Supplementary Figure 2.**
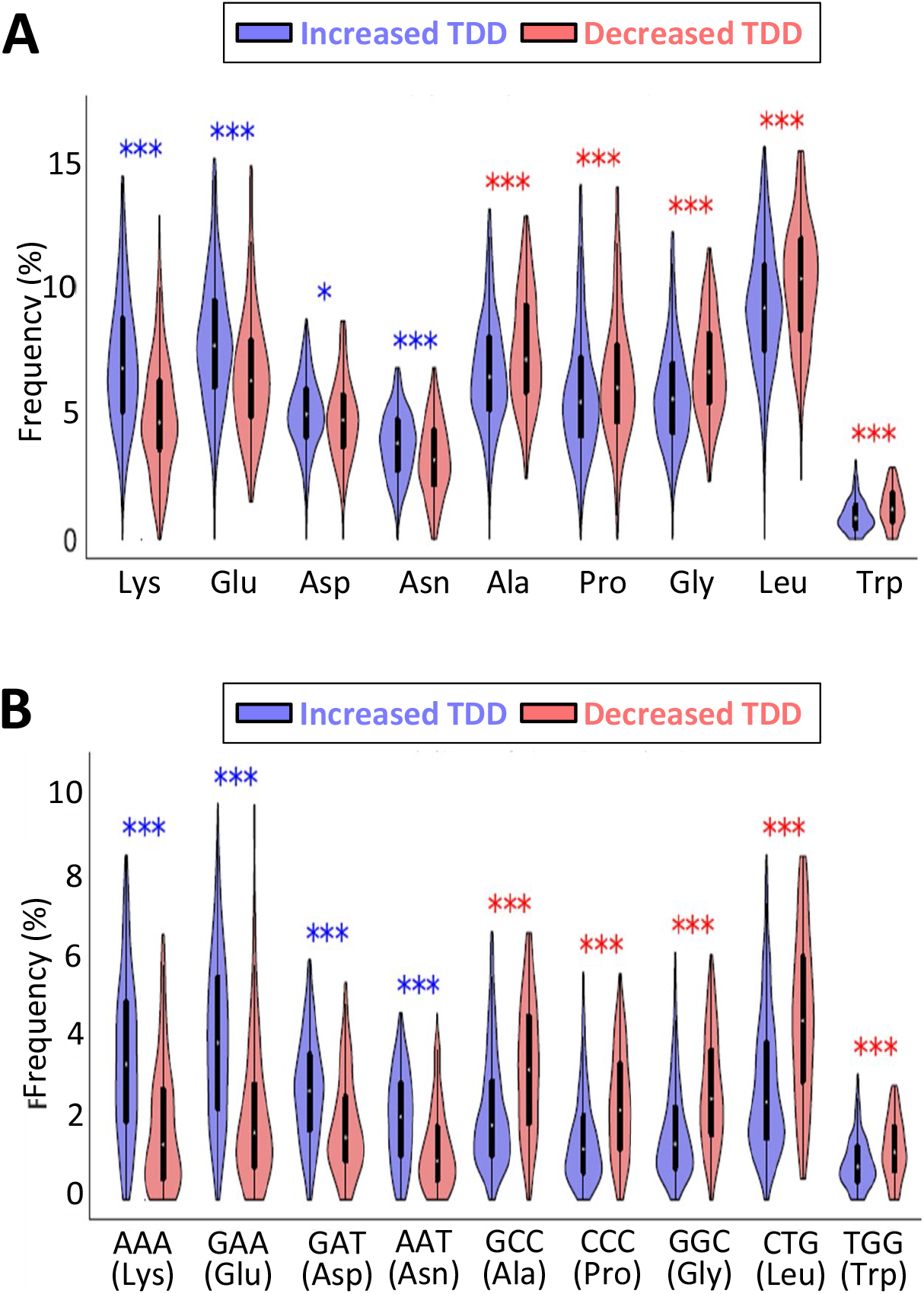
**a** Amino acid frequency in proteins encoded by mRNAs whose TDD index increased or decreased in response to MAPKi. *** and * mean that the frequency of an amino acid is statistically different (beta regression analysis followed by a Tukey’s test (pairwise comparison) FDR ≤ 0.05 (*), <0.001 (***)) between proteins encoded by mRNAs whose TDD increased and proteins encoded by mRNAs whose TDD decreased in response to MAPKi. **b** Codon frequency in mRNAs whose TDD index increased or decreased in response to MAPKi. *** means that codons are statistically different (beta regression analysis followed by a Tukey’s test (pairwise comparison) FDR <0.001) between mRNAs whose TDD increased and mRNAs whose TDD decreased in response to MAPKi.

**Supplementary Figure 3.**
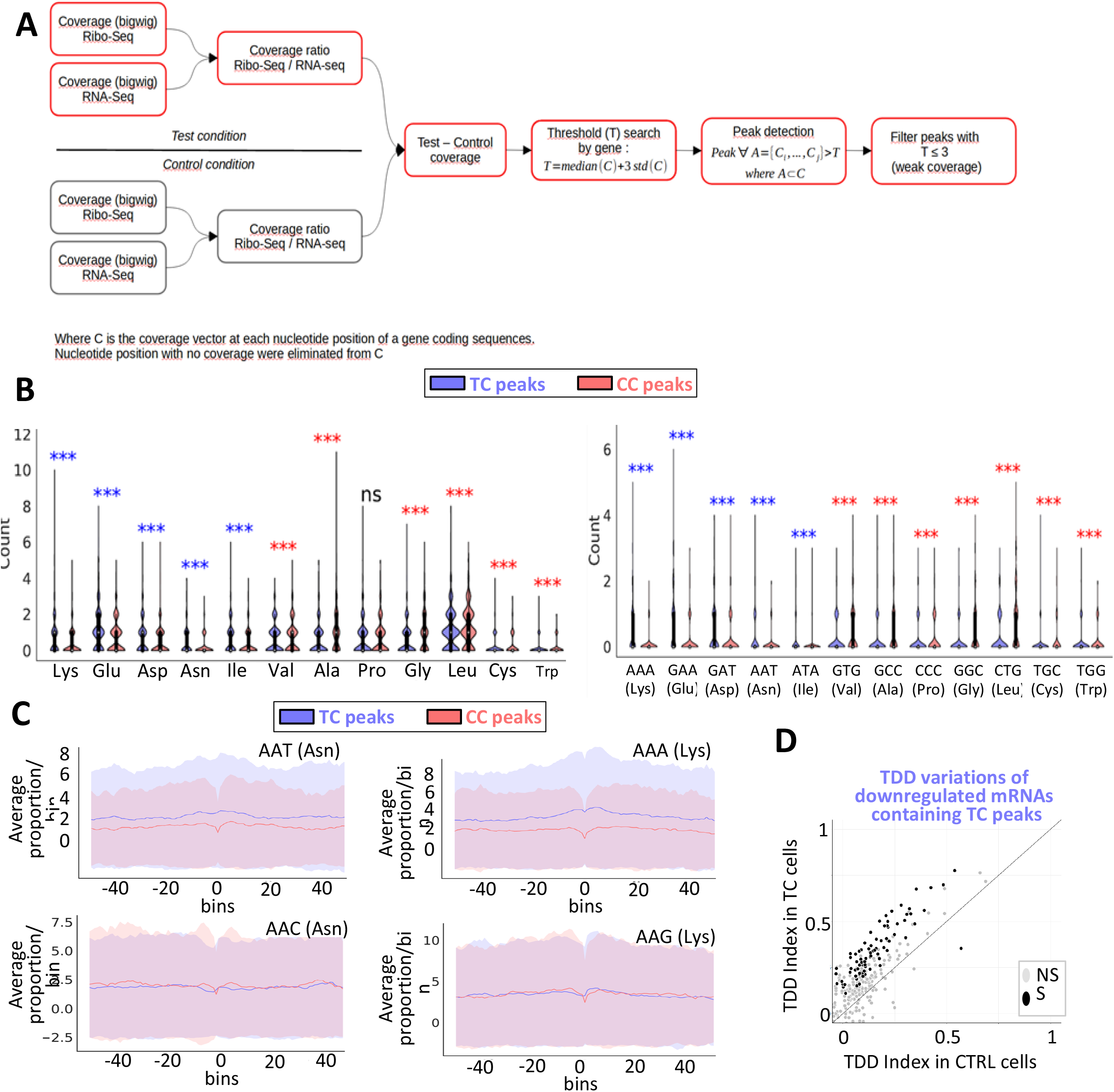
**a** Principle of the computational strategy to measure CC peaks and TC peaks (see Methods). **b** Amino acid (left panel) and codon (right panel) frequency in CC peaks and in TC peaks. *** means that codons or amino acids counts are statistically different (zero inflated negative binomial regression analysis FDR < 0.001) between TC peaks and in CC peaks. **c** Frequencies of codons within and around ribosome peaks. The average frequencies of codons at bin 0 was computed in ribosome protected mRNA regions. The same procedure was applied for other bins (windows of 10 codons) upstream and downstream starting from the central coordinate of each peak. The red curve corresponds to the values computed from CC peaks and the red shadow reflects the standard deviation of the values. The blue curve corresponds to the values computed from TC peaks and the blue shadow reflects the standard deviation of the values. **d** Comparison of the TDD index of mRNAs measured in control cells to the TDD index of mRNAs that are downregulated by MAPKi treatment and that contain ribosomal TC peaks. The TDD index measured in control cells of each MAPKi-downregulated mRNA whose TDD increased (x-axis) was plotted against the TDD index measured in MAPKi-treated cells (y-axis). Grey dots represent mRNAs whose TDD index was not statistically different (NS) when comparing treated cells to control cells. Black dots represent mRNAs whose TDD index was statistically different (S, linear regression analysis t-test p-value ≤ 0.05) when comparing treated cells to control cells. The gray line indicates when the TDD values are identical under the compared conditions.

**Supplementary Figure 4.**
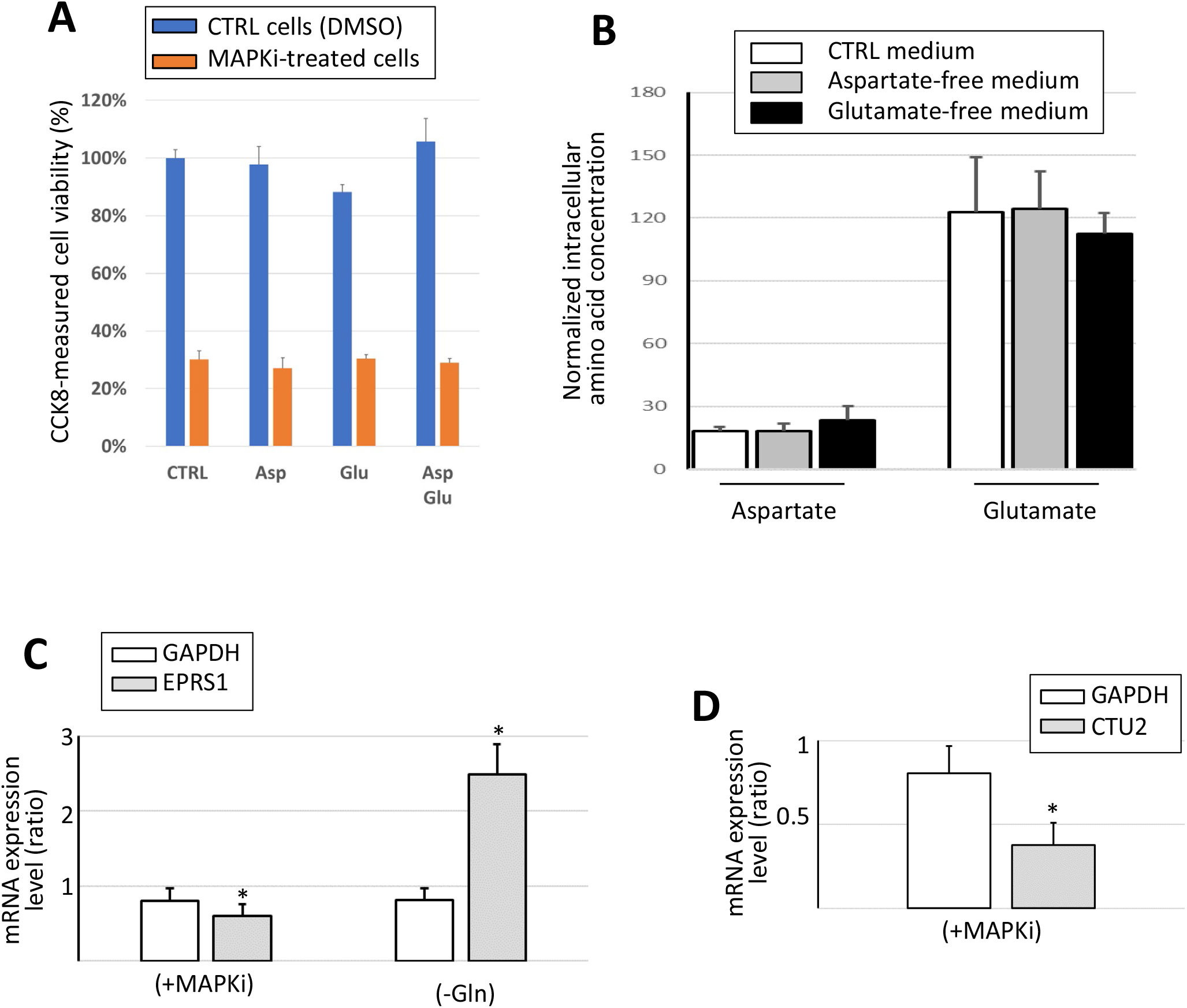
**a** Cell viability measured by the CCK8 assay after 24h of MAPKi treatment in a CTRL, Asp-free, Glu-free or (Asp+Glu)-free medium (n=3). **b** Asp and Glu intracellular concentration measured in cells grown for 24h in control or in aspartate-free or glutamate-free media (n=3). **c** RT-qPCR analysis of the expression level of the GAPDH and EPRS1 mRNAs in control cells, cells treated for 24h with MAPKi (MAPKi) (n=12), and cells grown for 24h in the absence of Gln (-Gln) (n=5). The values were normalized by the values obtained in control cells. * P≤0.05 (two-tailed paired t-test). **d** RT-qPCR analysis of the expression level of the GAPDH and CTU2 mRNAs in control cells and cells treated for 24h with MAPKi (+MAPKi) (n=12). The values were normalized by the values obtained in control cells. * P≤0.05 (two-tailed paired t-test).

## References

1. Wu, Q. & Bazzini, A.A. Translation and mRNA Stability Control. Annu Rev Biochem (2023).

2. Mishima, Y., Han, P., Ishibashi, K., Kimura, S. & Iwasaki, S. Ribosome slowdown triggers codon-mediated mRNA decay independently of ribosome quality control. EMBO J 41, e109256 (2022).

3. Bae, H. & Coller, J. Codon optimality-mediated mRNA degradation: Linking translational elongation to mRNA stability. Mol Cell 82, 1467–1476 (2022).

4. Morris, C., Cluet, D. & Ricci, E.P. Ribosome dynamics and mRNA turnover, a complex relationship under constant cellular scrutiny. Wiley Interdiscip Rev RNA 12, e1658 (2021).

5. Wu, Q. et al. Translation affects mRNA stability in a codon-dependent manner in human cells. Elife 8 (2019).

6. Bazzini, A.A. et al. Codon identity regulates mRNA stability and translation efficiency during the maternal-to-zygotic transition. EMBO J 35, 2087–2103 (2016).

7. Presnyak, V. et al. Codon optimality is a major determinant of mRNA stability. Cell 160, 1111–1124 (2015).

8. Peterson, J., Li, S., Kaltenbrun, E., Erdogan, O. & Counter, C.M. Expression of transgenes enriched in rare codons is enhanced by the MAPK pathway. Sci Rep 10, 22166 (2020).

9. Clarke, T.F.t. & Clark, P.L. Rare codons cluster. PLoS One 3, e3412 (2008).

10. Gillen, S.L., Waldron, J.A. & Bushell, M. Codon optimality in cancer. Oncogene 40, 6309–6320 (2021).

11. Hia, F. et al. Codon bias confers stability to human mRNAs. EMBO Rep 20, e48220 (2019).

12. Courel, M. et al. GC content shapes mRNA storage and decay in human cells. Elife 8 (2019).

13. Mauger, D.M. et al. mRNA structure regulates protein expression through changes in functional half-life. Proc Natl Acad Sci U S A 116, 24075–24083 (2019).

14. Nakamura, Y., Gojobori, T. & Ikemura, T. Codon usage tabulated from international DNA sequence databases: status for the year 2000. Nucleic Acids Res 28, 292 (2000).

15. Martin, S. et al. Oligodendrocyte differentiation alters tRNA modifications and codon optimality-mediated mRNA decay. Nat Commun 13, 5003 (2022).

16. Rezgui, V.A. et al. tRNA tKUUU, tQUUG, and tEUUC wobble position modifications fine-tune protein translation by promoting ribosome A-site binding. Proc Natl Acad Sci U S A 110, 12289–12294 (2013).

17. Rapino, F. et al. Wobble tRNA modification and hydrophilic amino acid patterns dictate protein fate. Nat Commun 12, 2170 (2021).

18. Rapino, F. et al. Codon-specific translation reprogramming promotes resistance to targeted therapy. Nature 558, 605–609 (2018).

19. Zhang, Z. et al. Global analysis of tRNA and translation factor expression reveals a dynamic landscape of translational regulation in human cancers. Commun Biol 1, 234 (2018).

20. Torrent, M., Chalancon, G., de Groot, N.S., Wuster, A. & Madan Babu, M. Cells alter their tRNA abundance to selectively regulate protein synthesis during stress conditions. Sci Signal 11 (2018).

21. Aharon-Hefetz, N. et al. Manipulation of the human tRNA pool reveals distinct tRNA sets that act in cellular proliferation or cell cycle arrest. Elife 9 (2020).

22. Gingold, H. et al. A dual program for translation regulation in cellular proliferation and differentiation. Cell 158, 1281–1292 (2014).

23. Thandapani, P. et al. Valine tRNA levels and availability regulate complex I assembly in leukaemia. Nature 601, 428–433 (2022).

24. Dittmar, K.A., Sorensen, M.A., Elf, J., Ehrenberg, M. & Pan, T. Selective charging of tRNA isoacceptors induced by amino-acid starvation. EMBO Rep 6, 151–157 (2005).

25. Passarelli, M.C. et al. Leucyl-tRNA synthetase is a tumour suppressor in breast cancer and regulates codon-dependent translation dynamics. Nat Cell Biol 24, 307–315 (2022).

26. Mazor, K.M. et al. Effects of single amino acid deficiency on mRNA translation are markedly different for methionine versus leucine. Sci Rep 8, 8076 (2018).

27. Darnell, A.M., Subramaniam, A.R. & O’Shea, E.K. Translational Control through Differential Ribosome Pausing during Amino Acid Limitation in Mammalian Cells. Mol Cell 71, 229–243 e211 (2018).

28. Loayza-Puch, F. et al. TGFbeta1-induced leucine limitation uncovered by differential ribosome codon reading. EMBO Rep 18, 549–557 (2017).

29. Loayza-Puch, F. et al. Tumour-specific proline vulnerability uncovered by differential ribosome codon reading. Nature 530, 490–494 (2016).

30. Shen, L. et al. SLC38A2 provides proline to fulfill unique synthetic demands arising during osteoblast differentiation and bone formation. Elife 11 (2022).

31. Kampen, K.R. et al. Translatome analysis reveals altered serine and glycine metabolism in T-cell acute lymphoblastic leukemia cells. Nat Commun 10, 2542 (2019).

32. Kay, E.J. et al. Cancer-associated fibroblasts require proline synthesis by PYCR1 for the deposition of pro-tumorigenic extracellular matrix. Nat Metab 4, 693–710 (2022).

33. Kay, E.J., Koulouras, G. & Zanivan, S. Regulation of Extracellular Matrix Production in Activated Fibroblasts: Roles of Amino Acid Metabolism in Collagen Synthesis. Front Oncol 11, 719922 (2021).

34. Eniafe, J. & Jiang, S. The functional roles of TCA cycle metabolites in cancer. Oncogene 40, 3351–3363 (2021).

35. Yoo, H.C., Yu, Y.C., Sung, Y. & Han, J.M. Glutamine reliance in cell metabolism. Exp Mol Med 52, 1496–1516 (2020).

36. Garcia-Bermudez, J. et al. Aspartate is a limiting metabolite for cancer cell proliferation under hypoxia and in tumours. Nat Cell Biol 20, 775–781 (2018).

37. Zhang, J., Pavlova, N.N. & Thompson, C.B. Cancer cell metabolism: the essential role of the nonessential amino acid, glutamine. EMBO J 36, 1302–1315 (2017).

38. Walker, M.C. & van der Donk, W.A. The many roles of glutamate in metabolism. J Ind Microbiol Biotechnol 43, 419–430 (2016).

39. Coloff, J.L. et al. Differential Glutamate Metabolism in Proliferating and Quiescent Mammary Epithelial Cells. Cell Metab 23, 867–880 (2016).

40. Pavlova, N.N., Zhu, J. & Thompson, C.B. The hallmarks of cancer metabolism: Still emerging. Cell Metab 34, 355–377 (2022).

41. Helenius, I.T., Madala, H.R. & Yeh, J.J. An Asp to Strike Out Cancer? Therapeutic Possibilities Arising from Aspartate’s Emerging Roles in Cell Proliferation and Survival. Biomolecules 11 (2021).

42. Melendez-Rodriguez, F. et al. HIF1alpha Suppresses Tumor Cell Proliferation through Inhibition of Aspartate Biosynthesis. Cell Rep 26, 2257–2265 e2254 (2019).

43. Sullivan, L.B. et al. Aspartate is an endogenous metabolic limitation for tumour growth. Nat Cell Biol 20, 782–788 (2018).

44. Yang, L., Venneti, S. & Nagrath, D. Glutaminolysis: A Hallmark of Cancer Metabolism. Annu Rev Biomed Eng 19, 163–194 (2017).

45. Ratnikov, B. et al. Glutamate and asparagine cataplerosis underlie glutamine addiction in melanoma. Oncotarget 6, 7379–7389 (2015).

46. Karki, P., Sensenbach, S., Angardi, V. & Orman, M.A. BRAF-Inhibitor-Induced Metabolic Alterations in A375 Melanoma Cells. Metabolites 11 (2021).

47. Karki, P., Angardi, V., Mier, J.C. & Orman, M.A. A Transient Metabolic State in Melanoma Persister Cells Mediated by Chemotherapeutic Treatments. Front Mol Biosci 8, 780192 (2021).

48. Alkaraki, A., McArthur, G.A., Sheppard, K.E. & Smith, L.K. Metabolic Plasticity in Melanoma Progression and Response to Oncogene Targeted Therapies. Cancers (Basel) 13 (2021).

49. Cesi, G., Walbrecq, G., Zimmer, A., Kreis, S. & Haan, C. ROS production induced by BRAF inhibitor treatment rewires metabolic processes affecting cell growth of melanoma cells. Mol Cancer 16, 102 (2017).

50. Corazao-Rozas, P. et al. Mitochondrial oxidative phosphorylation controls cancer cell’s life and death decisions upon exposure to MAPK inhibitors. Oncotarget 7, 39473–39485 (2016).

51. Haq, R., Fisher, D.E. & Widlund, H.R. Molecular pathways: BRAF induces bioenergetic adaptation by attenuating oxidative phosphorylation. Clin Cancer Res 20, 2257–2263 (2014).

52. Liang, S., Ezerskyte, M., Wang, J., Pelechano, V. & Dreij, K. Transcriptional mutagenesis dramatically alters genome-wide p53 transactivation landscape. Sci Rep 10, 13513 (2020).

53. Konovalov, K.A. et al. 8-Oxo-guanine DNA damage induces transcription errors by escaping two distinct fidelity control checkpoints of RNA polymerase II. J Biol Chem 294, 4924–4933 (2019).

54. Bradley, C.C., Gordon, A.J.E., Halliday, J.A. & Herman, C. Transcription fidelity: New paradigms in epigenetic inheritance, genome instability and disease. DNA Repair (Amst) 81, 102652 (2019).

55. Morreall, J.F., Petrova, L. & Doetsch, P.W. Transcriptional mutagenesis and its potential roles in the etiology of cancer and bacterial antibiotic resistance. J Cell Physiol 228, 2257–2261 (2013).

56. Bregeon, D. & Doetsch, P.W. Transcriptional mutagenesis: causes and involvement in tumour development. Nat Rev Cancer 11, 218–227 (2011).

57. Bregeon, D., Peignon, P.A. & Sarasin, A. Transcriptional mutagenesis induced by 8-oxoguanine in mammalian cells. PLoS Genet 5, e1000577 (2009).

58. Saxowsky, T.T., Meadows, K.L., Klungland, A. & Doetsch, P.W. 8-Oxoguanine-mediated transcriptional mutagenesis causes Ras activation in mammalian cells. Proc Natl Acad Sci U S A 105, 18877–18882 (2008).

59. Birnbaum, M.D., Nemzow, L., Kumar, A., Gong, F. & Zhang, F. A Rapid and Precise Mutation-Activated Fluorescence Reporter for Analyzing Acute Mutagenesis Frequency. Cell Chem Biol 25, 1038–1049 e1035 (2018).

60. Avagliano, A. et al. Metabolic Plasticity of Melanoma Cells and Their Crosstalk With Tumor Microenvironment. Front Oncol 10, 722 (2020).

61. Wasinger, C., Hofer, A., Spadiut, O. & Hohenegger, M. Amino Acid Signature in Human Melanoma Cell Lines from Different Disease Stages. Sci Rep 8, 6245 (2018).

62. Ratnikov, B.I., Scott, D.A., Osterman, A.L., Smith, J.W. & Ronai, Z.A. Metabolic rewiring in melanoma. Oncogene 36, 147–157 (2017).

63. Scott, D.A. et al. Comparative metabolic flux profiling of melanoma cell lines: beyond the Warburg effect. J Biol Chem 286, 42626–42634 (2011).

64. Afshar, N. et al. A novel motif of Rad51 serves as an interaction hub for recombination auxiliary factors. Elife 10 (2021).

65. Wang, H.C., Chou, C.C., Hsu, K.C., Lee, C.H. & Wang, A.H. New paradigm of functional regulation by DNA mimic proteins: Recent updates. IUBMB Life 71, 539–548 (2019).

66. Chou, C.C. & Wang, A.H. Structural D/E-rich repeats play multiple roles especially in gene regulation through DNA/RNA mimicry. Mol Biosyst 11, 2144–2151 (2015).

67. Hartwig, A. Role of magnesium in genomic stability. Mutat Res 475, 113–121 (2001).

68. Jia, D. et al. Drug-Tolerant Idling Melanoma Cells Exhibit Theory-Predicted Metabolic Low-Low Phenotype. Front Oncol 10, 1426 (2020).

69. Shen, S., Vagner, S. & Robert, C. Persistent Cancer Cells: The Deadly Survivors. Cell 183, 860–874 (2020).

70. Bristot, I.J., Kehl Dias, C., Chapola, H., Parsons, R.B. & Klamt, F. Metabolic rewiring in melanoma drug-resistant cells. Crit Rev Oncol Hematol 153, 102995 (2020).

71. Ramirez, M. et al. Diverse drug-resistance mechanisms can emerge from drug-tolerant cancer persister cells. Nat Commun 7, 10690 (2016).

72. Zhang, G. et al. Targeting mitochondrial biogenesis to overcome drug resistance to MAPK inhibitors. J Clin Invest 126, 1834–1856 (2016).

73. Baenke, F. et al. Resistance to BRAF inhibitors induces glutamine dependency in melanoma cells. Mol Oncol 10, 73–84 (2016).

74. Chen, S., Zhou, Y., Chen, Y. & Gu, J. fastp: an ultra-fast all-in-one FASTQ preprocessor. Bioinformatics 34, i884–i890 (2018).

75. Adiconis, X. et al. Comparative analysis of RNA sequencing methods for degraded or low-input samples. Nat Methods 10, 623–629 (2013).

76. Heyer, E.E., Ozadam, H., Ricci, E.P., Cenik, C. & Moore, M.J. An optimized kit-free method for making strand-specific deep sequencing libraries from RNA fragments. Nucleic Acids Res 43, e2 (2015).

77. Modolo, L. & Lerat, E. UrQt: an efficient software for the Unsupervised Quality trimming of NGS data. BMC Bioinformatics 16, 137 (2015).

78. Kim, D., Paggi, J.M., Park, C., Bennett, C. & Salzberg, S.L. Graph-based genome alignment and genotyping with HISAT2 and HISAT-genotype. Nat Biotechnol 37, 907–915 (2019).

79. Ramirez, F. et al. deepTools2: a next generation web server for deep-sequencing data analysis. Nucleic Acids Res 44, W160–165 (2016).

80. Sherman, B.T. et al. DAVID: a web server for functional enrichment analysis and functional annotation of gene lists (2021 update). Nucleic Acids Res 50, W216–W221 (2022).

